# Co-existence of synaptic plasticity and metastable dynamics in a spiking model of cortical circuits

**DOI:** 10.1101/2023.12.07.570692

**Authors:** Xiaoyu Yang, Giancarlo La Camera

## Abstract

Evidence for metastable dynamics and its role in brain function is emerging at a fast pace and is changing our understanding of neural coding by putting an emphasis on hidden states of transient activity. Clustered networks of spiking neurons have enhanced synaptic connections among groups of neurons forming structures called cell assemblies; such networks are capable of producing metastable dynamics that is in agreement with many experimental results. However, it is unclear how a clustered network structure producing metastable dynamics may emerge from a fully local plasticity rule, i.e., a plasticity rule where each synapse has only access to the activity of the neurons it connects (as opposed to the activity of other neurons or other synapses). Here, we propose a local plasticity rule producing ongoing metastable dynamics in a deterministic, recurrent network of spiking neurons. The metastable dynamics co-exists with ongoing plasticity and is the consequence of a self-tuning mechanism that keeps the synaptic weights close to the instability line where memories are spontaneously reactivated. In turn, the synaptic structure is stable to ongoing dynamics and random perturbations, yet it remains sufficiently plastic to remap sensory representations to encode new sets of stimuli. Both the plasticity rule and the metastable dynamics scale well with network size, with synaptic stability increasing with the number of neurons. Overall, our results show that it is possible to generate metastable dynamics over meaningful hidden states using a simple but biologically plausible plasticity rule which co-exists with ongoing neural dynamics.

## 1 Introduction

Cortical circuits express ongoing neural dynamics that is often found to be metastable, i.e., to unfold as a sequence of neural activity patterns in which ensembles of neurons keep approximately constant firing rates for transient periods of time. A stable vector of firing rates across simultaneously recorded neurons may be thought of as a ‘hidden state’, of which spike trains emitted by the neurons are noisy observations. In the cortex of rodents and non-human primates, metastable states last between a few hundred of ms to a few seconds and their identity and dynamics have been linked to sensory processes, attention, expectation, navigation, decision making and behavioral accuracy (see (La Camera et al., 2019; Brinkman et al., 2022) for reviews). One way to model this type of metastable activity is to organize a neural network in clusters, or cell assemblies (Litwin-Kumar and Doiron, 2012; Mazzucato et al., 2015). The neurons in each clusters are connected by synaptic weights whose average value is larger than the average weight between neurons of different clusters. A key question is then to understand how such structure subserving metastable dynamics can emerge from, for example, experience-dependent plasticity. This problem of structuring a neural circuit in clusters via synaptic plasticity is basically the same problem of the stable formation of Hebb assemblies leading to persistent activity reflecting the memory of learned stimuli (Del Giudice et al., 2003; Amit and Mongillo, 2003; Litwin-Kumar and Doiron, 2014; Zenke et al., 2015).

The problem of the formation of stable cell assemblies has been of interest for a long time (see (Dayan and Abbott, 2005; Gerstner et al., 2014) for textbook reviews). The most recent efforts in this direction have included the combination of spike-timing-dependent plasticity (STDP) (Markram et al., 1997; Bi and Poo, 1998) and a number of homeostatic mechanisms (Watt and Desai, 2010; Turrigiano, 2017) to keep the neural activity bounded during learning. However, while most efforts so far have focused on the formation of stable neural clusters with the purpose of representing retrievable memories or the development of receptive fields, here we focus on metastable dynamics. In other words, instead of focusing on stable neural dynamics following the presentation and removal of a stimulus, the aim of this study is to obtain neural dynamics that continuously switches among a set of hidden states, which have been stored in the network structure by training. The ensuing switching dynamics can be interpreted as a continuous reactivation of internal representations. Aside from potential computational consequences, we motivate our quest from the observation of switching dynamics in many neuroscience studies (reviewed in (La Camera et al., 2019; Brinkman et al., 2022)).

In pursuing this effort, we require the synaptic plasticity rule to be biologically plausible and leading to the formation of neural clusters that are stable for random perturbations. Moreover, we require that the neural clusters generate metastable dynamics that coexists with ongoing plasticity (i.e., in the absence of external stimuli). We further aim to obtain a model where new information can be accommodated, so that training with a new set of stimuli will lead to cluster rearrangement producing metastable dynamics among the new states. The requirement of biological plausibility mostly means that the plasticity rule must be local and must depend only on presynaptic and postsynaptic activity in a way that is accessible to the synapse – in particular, it must depend only on presynaptic spikes and postsynaptic variables related e.g. to membrane voltage or calcium transients (Fusi et al., 2000; Shouval et al., 2002; Clopath et al., 2010; Graupner and Brunel, 2012).

To our knowledge, one previous study has provided a model of synaptic plasticity that can produce metastable dynamics, seen as a signature of slow fluctuations in neural circuits (Litwin-Kumar and Doiron, 2014). This model uses a combination of STDP and inhibitory plasticity, plus a non-local mechanism of synaptic renormalization. In contrast, here we present a plasticity rule that only on presynaptic spikes, postsynaptic membrane potential and postsynaptic spikes. Long-term potentiation (LTP) and long-term depression (LTD) are obtained by comparing a voltage-sensitive internal variable to an adapting threshold, reminiscent of the BCM rule (Bienenstock et al., 1982). The threshold adapts in a way to produce transient LTP among co-active neurons, followed by LTD after prolonged activation. This leads to the stable formation of neural clusters while promoting a dynamics that is automatically metastable for a large range of network sizes. The learning rule keeps the synaptic weights near a critical line where metastable dynamics is the only equilibrium dynamics of the network, as the result of a self-tuning mechanism that keeps the network activity near the threshold for memory reactivation. This is accomplished without additional homeostatic mechanisms, such as inhibitory plasticity or synaptic scaling (Tetzlaff et al., 2011; Zenke and Gerstner, 2017; Weidel et al., 2021), streamlining the plasticity rule to require only a handful of basic ingredients.

In summary, our plasticity rule thus provides a possible explanation of the emergence of metastable dynamics observed in many brain areas during both ongoing and evoked neural activity, and provides a self-tuning mechanism that keeps the network activity near a critical line characterized by slow fluctuations and spontaneous memory reactivations.

## 2 Results

### 2.1 The synaptic plasticity rule

We endowed a recurrent network of excitatory and inhibitory exponential integrate-and-fire (EIF) neurons (see Methods) with the following plasticity rule for synapses connecting excitatory neurons. Given a synapse with efficacy *w*_*ij*_ from excitatory neuron *j* to excitatory neuron *i*, a change in synaptic efficacy was triggered by the arrival of a presynaptic spike, while its polarity and strength depended on the activity of the postsynaptic neuron:

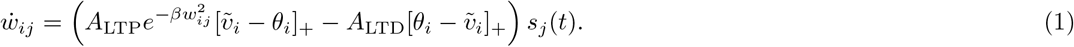

Here,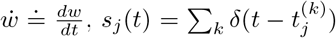 is the presynaptic spike train and *δ*(*t*) is the Dirac delta function. [*x*]_+_ ≐ max(*x*, 0) is the rectified linear function and 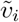 is a local postsynaptic variable which dictates the polarity of plasticity: the synapse undergoes LTP when 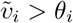, and it undergoes LTD when 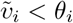. To respect Dale’s law, the weights are constrained to be nonnegative.

The variable 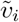 could represent a calcium variable or the running average of the membrane potential. In our case, 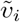 was the low-pass filter of 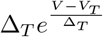, where *V* is the membrane potential of the EIF neuron (see Methods), and therefore it was driven most substantially during the emission of a spike. The strength of LTP was further modulated by an attenuation factor 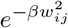 that constrains the ability of strong synapses to grow excessively.

As similar learning rules of this kind (Dayan and Abbott, 2005), this rule is unstable if *θ*_*i*_ is a constant threshold and the attenuation factor is missing. In such a case, stimulating a subset of neurons would result in higher firing rates and larger 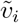, which in turn would lead to higher firing rates, and so on. One way to prevent this problem is to use activity-dependent thresholds as in the BCM rule (Bienenstock et al., 1982). This idea requires the LTP and LTD thresholds to be a non-linear function of the postsynaptic activity 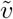, with faster dynamics than the dynamics of the synaptic weights. While the BCM rule uses a supralinear function of 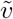, we chose a hyperbolic tangent function of both 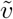 and postsynaptic spiking activity, 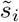:

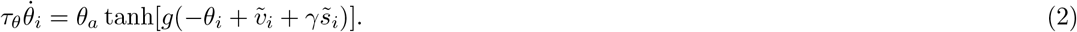

In this equation, 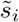 is the low-pass filtered postsynaptic spike train, 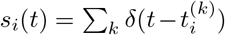, *θ*_*a*_ is a constant in units of *θ*_*i*_, *g* is a gain factor and *γ* is a constant. We use here the hyperbolic tangent function for convenience, however the exact form of the sigmoidal function is not essential.

The hyperbolic tangent in Eq. 2 automatically adjusts the dynamics of the threshold *θ*_*i*_(*t*) for different postsynaptic activities. The motivation behind this specific model is that it can lead naturally to switching dynamics of neural clusters. This can be understood from the dynamics of Eq. 2, which adapts to the size of its argument 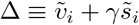 (Fig. 1a; see Methods for details). For small arguments, the dynamics is fastest and *θ*_*i*_ closely follows 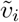, resulting in no average change in the synaptic weights (Fig. 1b). This occurs when the postsynaptic membrane potential is characterized by subthreshold fluctuations. When the postsynaptic neuron fires a spike, its membrane potential rises significantly and *θ*_*i*_ is attracted to a new value with slower dynamics. As a result, *θ*_*i*_ will temporarily lag behind 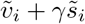, producing a short temporal window for LTP, followed by a longer window for LTD (Fig. 1c). An occasional spike during ongoing activity will not produce a meaningful synaptic change, however, repeated activation of the same postsynaptic neuron will produce a longer window for LTP (Fig. 1d, blue shaded area). This will occur when the postsynaptic neuron engages in recurrent excitation, and will promote the formation of clusters of co-active neurons. Prolonged activation of the same cluster, however, will turn LTP into LTD. This is due to the term 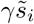, so that the threshold *θ*_*i*_ will finally approach a value 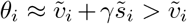, causing LTD (Fig. 1d, red shaded area). In summary, *θ*_*i*_ dynamics can help to form neural clusters via transient LTP, while capping the growth of synaptic efficacies via LTP to LTD transitions. Later we show that, upon repeated presentation of external inputs, our learning rule builds a synaptic structure that supports an equilibrium switching dynamics among co-active groups of neurons.

**Figure 1:**
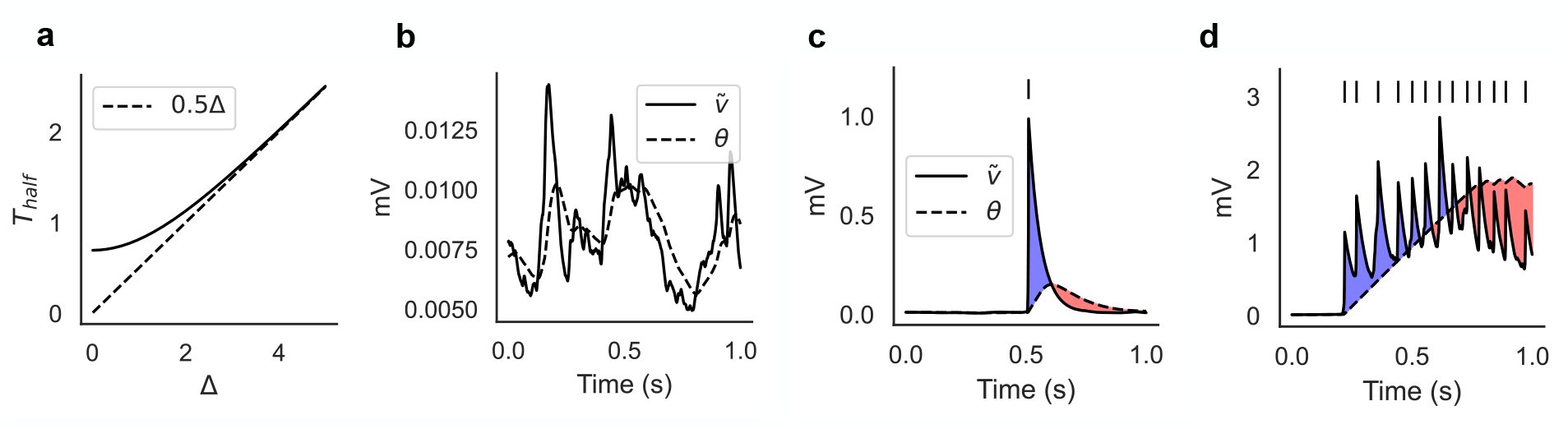
Illustration of the plasticity rule. (a) Time scale of 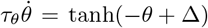 as a function of Δ, where 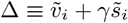 (see Eq. 2). *T*_half_ is defined as the time it takes for *θ*(0) = 0 to reach half of Δ (in units of *τ*_*θ*_), and for Δ ≳ 3 it increases linearly with Δ (see Methods). (b) The learning rule ignores the subthreshold fluctuations of the membrane potential when the neuron does not fire spikes. (c) After the postsynaptic neuron fires a single spike, the synapse undergoes a short window for LTP (blue shaded area) followed by a longer window for LTD (red shaded area). (d) Repeated activation of the postsynaptic neuron will cause a transition from LTP to LTD. Panels *c* and *d* are illustrative cartoons.

Note that the additive form 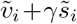 makes it easy to control plasticity for two different cases, one when the postsynaptic neuron is inactive (in which case 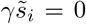, *θ*_*i*_ quickly follows 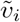, and no net plasticity ensues), and one during repetitive firing (in which case first LTP, and then LTD, ensues).

### 2.2 Formation of stable clusters with metastable dynamics

We initially tested our plasticity rule in a recurrent spiking network model comprising *N*_*E*_ = 800 excitatory and *N*_*I*_ = 200 inhibitory EIF neurons (see Methods). During training, a set of *Q* = 10 sensory stimuli were presented to the network in random order, each targeting a fixed but randomly chosen subpopulation of excitatory neurons, which we call a cluster. Although each cluster was associated to a separate stimulus, each cluster contained *f* = 10% of randomly selected neurons, so that the same neuron could respond to more than one stimulus as typically observed in experiments (Miyashita and Chang, 1988; Miyashita, 1988; Curti et al., 2004; Jezzini et al., 2013; Fusi et al., 2016). Thus, the mean number of neurons in each cluster was 80, but each neuron had a probability of about 0.26 to respond to at least two stimuli, and clusters had a probability 0.61 of sharing neurons with other clusters (see Methods for details).

Stimulus presentations occurred every 2000 ms and each lasted 500 ms (however, using random inter-stimulus intervals did not alter the results). This training procedure lasted for 10 minutes and on average each sensory stimulus was presented 30 times.

As expected, after training the network exhibited metastable ongoing dynamics which was still present after 2 and 4 hours (Fig. 2a). In this figure, spikes from neurons in the same cluster have the same color, and neurons belonging to multiple clusters are duplicated. The black curves superimposed to each cluster’s spikes measure the ‘overlap’ of the whole network’s activity with the stimulus associated to that cluster (see Methods). For a given stimulus, the overlap varies between zero and one and approaches 1 when all active excitatory neurons of the network belong to the associated cluster, while the remaining neurons have negligible firing rate. The raster plots in Fig. 2 show that the neural activity after training switches between networks states characterized by specific cluster activations. These states can therefore be interpreted as memories of the stimuli, with the ongoing dynamics being akin to a random walk among these memory states. The distribution of state durations was approximately exponential with mean around 250 − 300 ms (Fig. 2c), reminiscent of the discrete Markov processes with fast state transitions found to describe ensembles of cortical spike trains (Seidemann et al., 1996; Jones et al., 2007; Mazzucato et al., 2015; Maboudi et al., 2018; Benozzo et al., 2021; Lang et al., 2023). The metastable dynamics observed after training is the consequence of potentiated synapses inside clusters and the emergence of block structure in the synaptic matrix (Fig. 2b), a structure known to potentially produce metastable dynamics in spiking networks (Litwin-Kumar and Doiron, 2012; Deco and Hugues, 2012; Mazzucato et al., 2015). We present a more detailed analysis of this behavior in Sec. 2.5.

**Figure 2:**
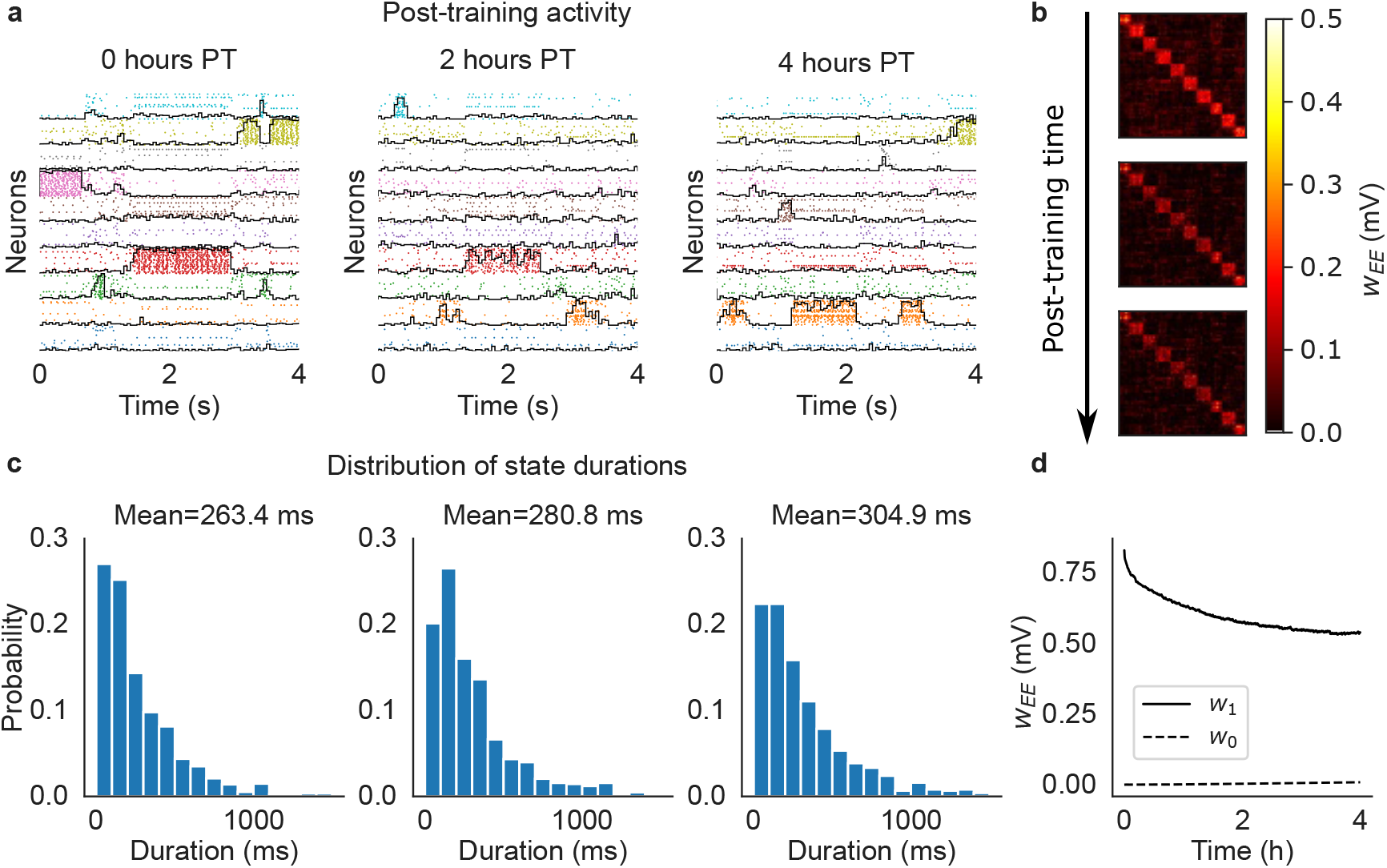
Cluster formation and metastable dynamics via excitatory plasticity in a network with 1, 000 spiking neurons (see the text). (a) Rasterplot of excitatory neurons taken immediately after training (left), 2 hours after training (middle), and 4 hours after training (right). Neurons were ordered according to cluster membership (different colors for different clusters), and if necessary also duplicated to multiple clusters according to the sensory stimuli they respond to. *Black lines*: overlap between the network’s neural activity and the stimulus associated to each cluster (see the text). (b) Synaptic matrix of the network at the same times as in (a) showing the formation of clusters (evidence from the block structure of the matrix). (c) Distribution of durations of cluster activations (see the text) for each corresponding epoch shown in (a). Samples were taken over a period of 10 minutes. (d) Averaged post-training excitatory synaptic weights as a function of time. *w*_1_: mean weights across synapses connecting neurons sharing at least one stimulus; *w*_0_: mean weights across synapses connecting neurons sharing no stimuli.

We next illustrate the role of the attenuation factor 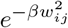 in Eq. 1 and the spiking term 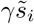 in Eq. 2. Without attenuation, training is successful, however clusters are unstable and disappear within one hour after training (Fig S1). Without the term 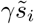 in Eq. 2, training produces uneven clusters that are unable to sustain cluster activation within one hour after training (Fig S2).

To quantify the amount of learning, we measured the average synaptic weight among synapses connecting neurons sharing at least one sensory stimulus (dubbed ‘*w*_1_’), and the average weight among neurons that did not share any sensory stimuli (‘*w*_0_’):

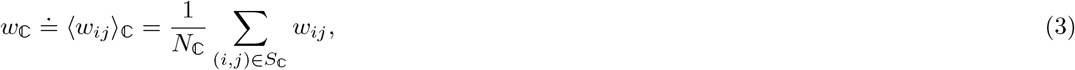

where ℂ ∈ {0, 1}, *N*_*ℂ*_ is the number of synapses of type ℂ, and *S*_*ℂ*_ is the set of *ij* indices of synapses of type ℂ. Training significantly increased *w*_1_ compared to *w*_0_, thus mapping successfully the association between sensory inputs and the corresponding neural clusters activated by the stimuli. After training, the excitatory synapses were continuously modified by the plasticity rule during ongoing metastable activity, leading to the decrease of *w*_1_ and increase of *w*_0_ (Fig. 2d). These synaptic changes seem to stabilize after 4 hours; after this time, plasticity coexists with metastable dynamics, the latter unfolding as a random walk among states associated with the training stimuli. This picture is confirmed in longer simulations of larger networks presented in later sections.

### 2.3 Robustness to perturbations and remapping

In the previous section we have shown that our learning rule generates metastable dynamics that co-exists with synaptic plasticity. We show next that this is also true in the presence of random stimuli. After training, we probed the network with external sensory inputs with similar physical features (i.e. magnitude and coding level) as the training stimuli. The occurrence schedule of these external stimuli was modeled as a Poisson process with a mean inter-event interval of 10 s. We considered three different scenarios. In the first scenario, the external stimuli were randomly sampled and used only once (i.e., the subsets of target neurons were uniformly and independently sampled at each occurrence; Fig. 3a). This scenario mimics a purely noisy environment where there is no temporal correlation in the sensory inputs. In the second scenario, only half the sensory inputs were random inputs, while the other half were sampled from a finite set of 10 sensory stimuli targeting always the same neurons (Fig. 3b). This scenario mimics a combination of meaningful stimuli occurring amid some random sensory background. Finally, in the third scenario the external stimuli were all sampled from a pre-defined set of 10 sensory stimuli, mimicking a situation where the network is being retrained with new stimuli (Fig. 3c). In all cases, the network kept adjusting its synaptic weights according to our plasticity rule.

**Figure 3:**
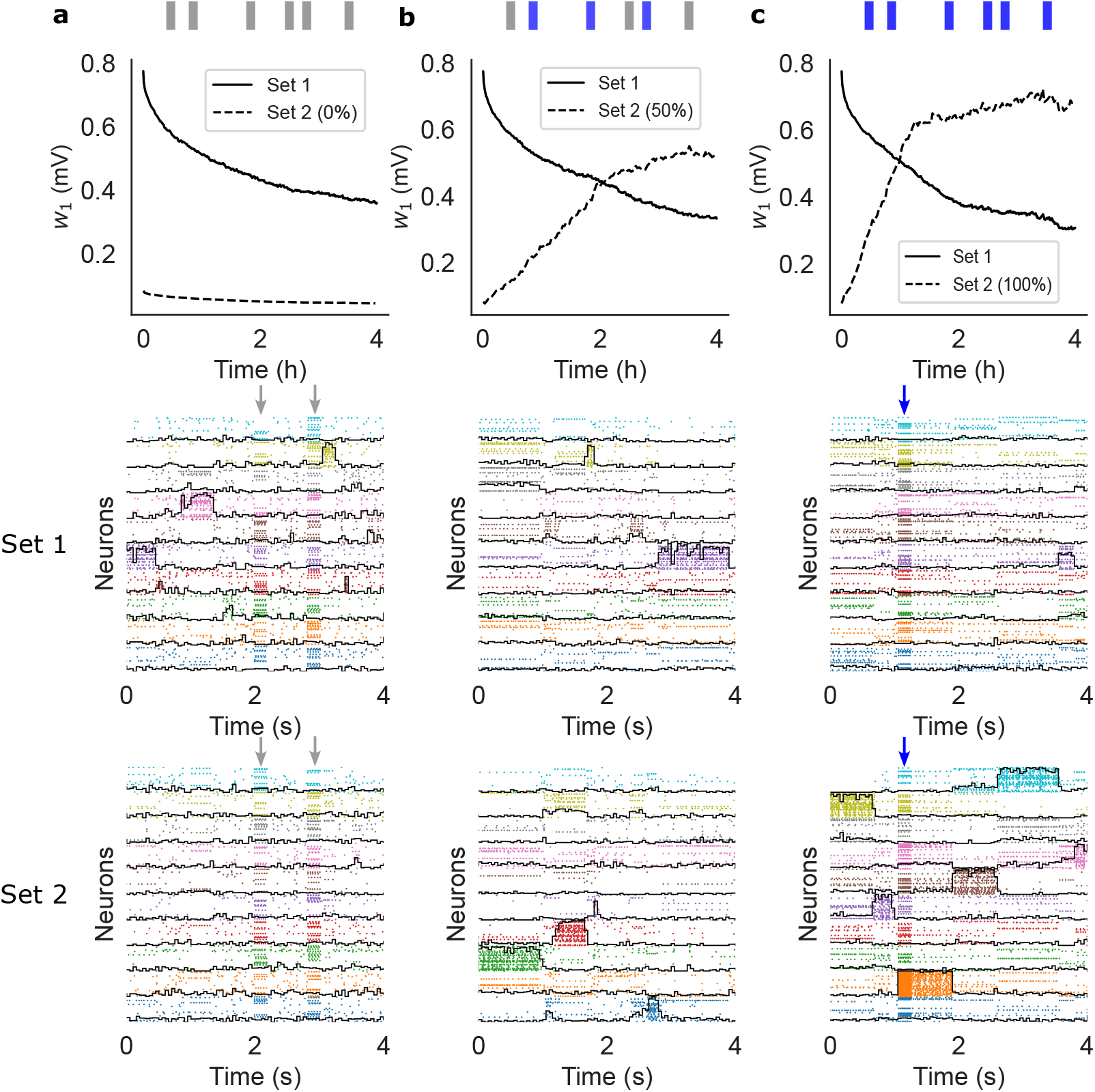
Sensory perturbations and remapping of the network after training (same network as Fig. 2). The network was trained with stimuli of Set 1 and then probed for after training, stimuli were drawn from a different set (Set 2) and delivered at random times. (a) Perturbing stimuli were randomly sampled to mimic a noisy environment. (b) 50% of perturbing stimuli were sampled randomly while the other 50% were sampled from a finite pre-defined set of reoccurring stimuli. (c) All stimuli were sampled from a finite pre-defined set of reoccurring stimuli. The top panels show *w*_1_ (Eq. 3) after training for Set 1 (full) and Set 2 (dashed). The middle panels show raster plots of the neural activity 4 hours after training, together with the overlaps (black lines) with the stimuli of Set 1 (those used for training). The bottom panels show raster plots of the neural activity 4 hours after training, together with the overlaps with the stimuli of Set 2 (note that this set was never used in (a)). Vertical arrows indicate an external stimulation with either a random stimulus (grey arrows) or a stimulus from Set 2 (blue arrows).

In the presence of random stimulation, the network maintained a significant memory of the learned stimuli for several hours, similar to what we found in the absence of sensory stimulation (Fig. 3a, top and middle panels). In the third scenario, the network quickly learned the new sensory inputs (Fig. 3c-bottom) while gradually forgetting the previously learned ones (Fig. 3c-middle). Finally, in the ‘mixed’ stimulation scenario, the network could still learn the new sensory stimuli but at a much slower rate (Fig. 3b-top, compare dashed lines in Figures 3b-top and 3c-top). A raster plot of the neural activity shows the metastable activation of clusters of neurons representing stimuli in both the first and second set (Fig. 3b, middle and bottom panels, respectively), showing that the network could accommodate the learning of new stimuli (Fig. 3b, bottom) while maintaining a trace of the previously learned ones. Note how, in between stimulus presentations, the network dynamics was metastable in all cases.

### 2.4 Learning stability vs. network size

Although synaptic weights decay after training (Fig. 2d), here we show that such decay slows down as a function of network size. We show this by estimating the rate of change for *w*_0_ and *w*_1_ as a function of N, the number of neurons in the network. Scaling up the network size can be done in several ways (Renart et al., 2007; Doiron and Litwin-Kumar, 2014); we used two different scaling procedures, one in which *Q* ∝ *N* and one in which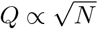.

In the first scenario, we scale up the number of clusters Q proportionally to *N*_*E*_ = 0.8N while keeping constant the mean cluster size *N*_*E*_/*Q* = 80 (this corresponds to a coding level *f* = 1/*Q* ∝ 1/*N*, where *f* is the probability that a neuron is targeted by a stimulus; see Methods). As shown in Fig. 4, synaptic decay is slower in larger networks (Fig. 4a) while ongoing metastable dynamics co-exists with synaptic plasticity (Fig. 4b). In particular, nearly constant values of the mean synaptic weights are observed in networks of 20, 000 neurons (rightmost panel of Fig. 4a).

**Figure 4:**
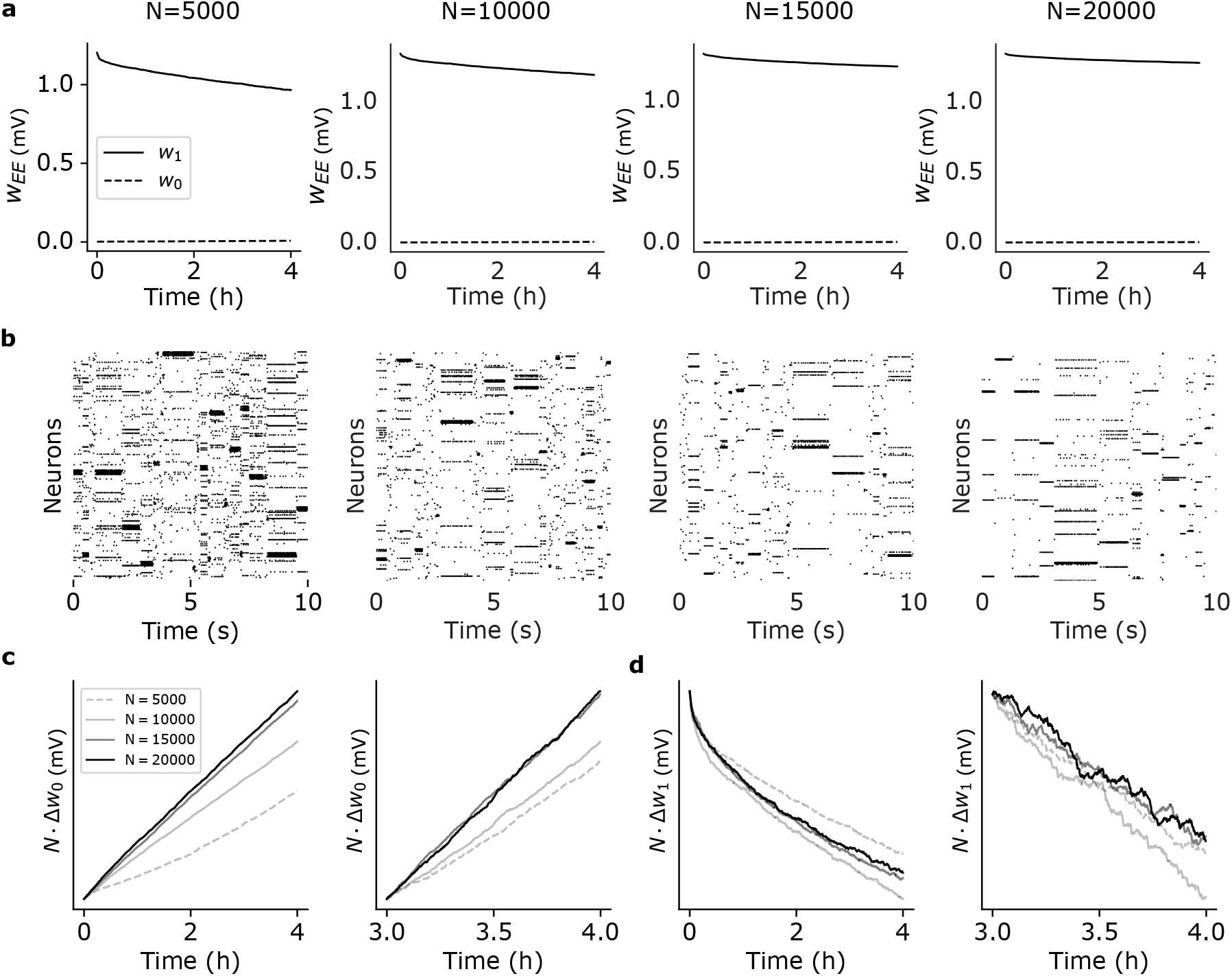
Effect of network size on synaptic dynamics after training. (a) Time dependence of average synaptic weights *w*_1_ and *w*_0_ after training for networks of different size *N* (same keys as Fig. 2d). In each case, the network had *N*_*E*_ = 0.8*N* excitatory neurons and *Q* = *N*_*E*_*/*80 = *N/*100 clusters (i.e., *Q* = 50, 100, 150, and 200 from left to right). (b) Raster plots of the network’s activity 4 hours after training for the corresponding networks in (a). (c) Plots of *N* Δ*w*_0_ vs. time after training for the different network in (a). Observations were taken 0 to 4 hours post training (left panel) and 3 to 4 hours post training (right panel). In large networks, *N* Δ*w*_0_ does not depend on *N* as predicted by Eq. 4. (d) Same as (c) for *N* Δ*w*_1_ vs. time.

Analytical arguments imply a synaptic decay rate ∝ *N* ^*−*1^, i.e.,

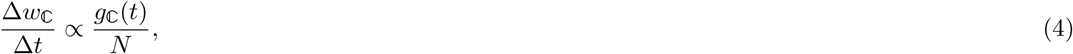

where Δ*w*_*ℂ*_ is the average change in strength over the interval Δ*t* for synapses of type ℂ (defined analogously to Eq. 3; see Methods) and *g*_*ℂ*_ (*t*) is a function of time but not of N. To confirm this prediction, we plotted *N*Δ*w*_0_ (Fig. 4c) and *N*Δ*w*_1_ (Fig. 4d) as a function of time and for increasing network size (from *N* = 5, 000 to *N* = 20, 000). The plots show that, after entering the ongoing metastable regime, both curves *N*Δ*w*_0_ and *N*Δ*w*_1_ tend to overlap for large enough networks, as predicted by Eq. 4. Empirically, *g*_*ℂ*_(*t*) is an increasing function for *w*_0_ synapses and a decreasing function for *w*_1_ synapses. The initial transients visible in the left panels of Fig. 4c-d are due to either a small *N* or not having yet reached the stable regime of synaptic decay (which takes about 4h in small networks, see Fig. 2d). For large *N* and starting 3h post training, the rate of change is independent of *N* (Fig. 4c-d, right panels).

These results confirm the synaptic decay rate ∝ *N* ^*−*1^ for *w*_1_ synapses, implying slower memory decay and more stable synaptic efficacies in larger networks, despite ongoing plasticity. Moreover, the dynamics produced by the networks of Fig. 4 shows near-exponential distributions of the state durations (Fig. S3), with mean durations approaching stability 4 hours post-training and for *N* > 15, 000.

The slowdown of synaptic decay rate with N was found also in an alternative scaling scenario in which *f* = 1/*Q* with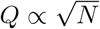, giving 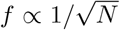 and 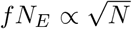 neurons in each cluster (with *N*_*E*_ = 0.8*N*).

In this case we have a smaller number of clusters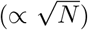, but each cluster grows in size with *N*. We trained several networks under this scaling and found results analogous to those of Fig. 4: training is successful, the synapses become more stable in larger networks, and mestastable dynamics is reliable across network sizes (see Fig S4). We could not estimate the synaptic rate of change in this case (see Methods). Empirically, the 1/*N* law was not observed in this case (Fig. S4c-d).

### 2.5 Mechanistic origin of neural metastability

What is the mechanism behind the the coexistence of synaptic plasticity and metastable dynamics? Due to the adaptive threshold in the learning rule, prolonged cluster activations are eventually terminated by LTD, however it is not clear what would cause the clusters’ re-activations ensuring an ongoing metastable regime. If, however, the synaptic structure reached by training satisfies some known criteria (Mazzucato et al., 2015), metastable dynamics would emerge due to endogenously generated fluctuations in the spiking network, aided by quenched random connectivity, sufficient synaptic potentiation inside clusters, and recurrent inhibition. In such a case, metastable dynamics would be present at the end of training also in the absence of plasticity. To show that this is indeed the case, we switched off synaptic plasticity at various times post-training and observed the neural dynamics. Specifically, we ran the network for 24 hours after training in the presence of synaptic plasticity and stored the synaptic matrix at 0, 1, 8 and 24 hours post-training. For each time point, we performed network simulations using the corresponding stored synaptic matrix, both with and without plasticity, for 10 minutes. The results are shown in Fig. 5 for a network of *N* = 5, 000 neurons and *Q* = 50 stimuli targeting non-overlapping clusters of neurons. Each row of the figure shows the synaptic weight distributions at a specific time point (left-most column), a snapshot of neural activity with (second column) and without (third column) ongoing plasticity, and normalized histograms of state durations with and without plasticity (right-most column). As shown in the figure, ongoing plasticity is not required for metastable dynamics, confirming that the latter is the consequence of endogenous fluctuations of the neural activity. The only appreciable difference between the two cases (plasticity ON vs. plasticity OFF) is seen 0 hours post-training: although metastable dynamics is present with or without plasticity, the histograms of state durations are different. This shows that, right after training, synaptic plasticity is *not necessary* for metastability, but it affects its statistics. One hour later (and in the following time points) the neural activities (and associated histograms) in the presence or absence of plasticity are indistinguishable (differences in mean state durations are the result of random fluctuations). This suggests that the synaptic weights were still converging, at time 0 post-training, towards a more stable region. In this latter region, metastable neural dynamics and synaptic dynamics coexist and generate the same neural activity that would be observed in the absence of plasticity. In such a phase, we expect the fluctuations in the synaptic weights caused by plasticity not to play a role in metastable dynamics.

**Figure 5:**
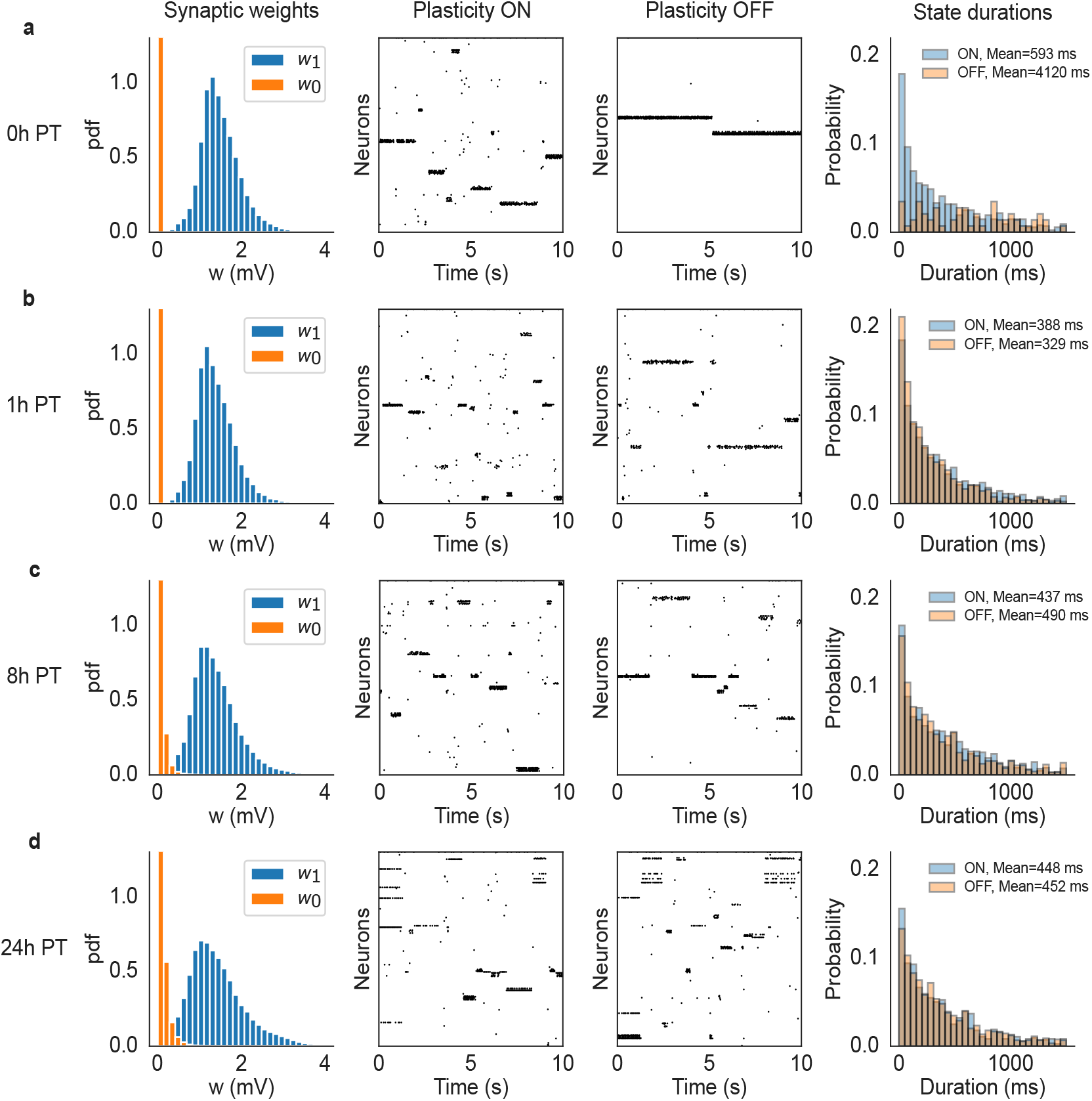
Comparison of network dynamics with and without synaptic plasticity in a network with *N* = 5000 neurons and *Q* = 50 stimuli. After training, the ongoing plasticity continued for 24 hours and we recorded the synaptic matrix at 0 (a), 1 (b), 8 (c) and 24 (d) hours post-training. Then we ran the network dynamics, starting with the corresponding stored synaptic matrix, for 10 minutes with and without plasticity respectively. Each row of the figure shows the synaptic weight distributions at each specific time point (left-most column), a snapshot of neural activity with (second column) and without (third column) ongoing plasticity, and histograms of state durations with and without plasticity (right-most column). The mean state durations are reported above the histograms.

To confirm this prediction, we performed a mean field analysis of the neural activity (see Methods). The analysis assumes, for simplicity, that only one cluster can be active at any given time. Above a critical value for the average synaptic value inside a cluster, metastable dynamics is possible and is revealed by a difference in firing rate between the active and non-active clusters (Mazzucato et al., 2015).

Fig. 6 (left panel) shows the mean field landscape of the neural activity. The landscape shows the mean-field predictions of the firing rate of the active cluster as a function of the mean and standard deviation of the synaptic weights inside the active cluster. The actual firing rates observed in the network are shown in the right panel. The contour lines shown in the figure are lines of equal firing rate. Below the lowest contour (firing rate ∼ 5 spikes/s), there is no predicted difference between the firing rates of active and inactive clusters, and the network is characterized by a spontaneous, low firing rate solution. Higher contour lines correspond to robust structuring of the synaptic weights, where the network is able to sustain persistent activity of its clusters (Amit and Brunel, 1997). With our learning rule, these higher lines are not reached due to LTP→LTD transitions.

**Figure 6:**
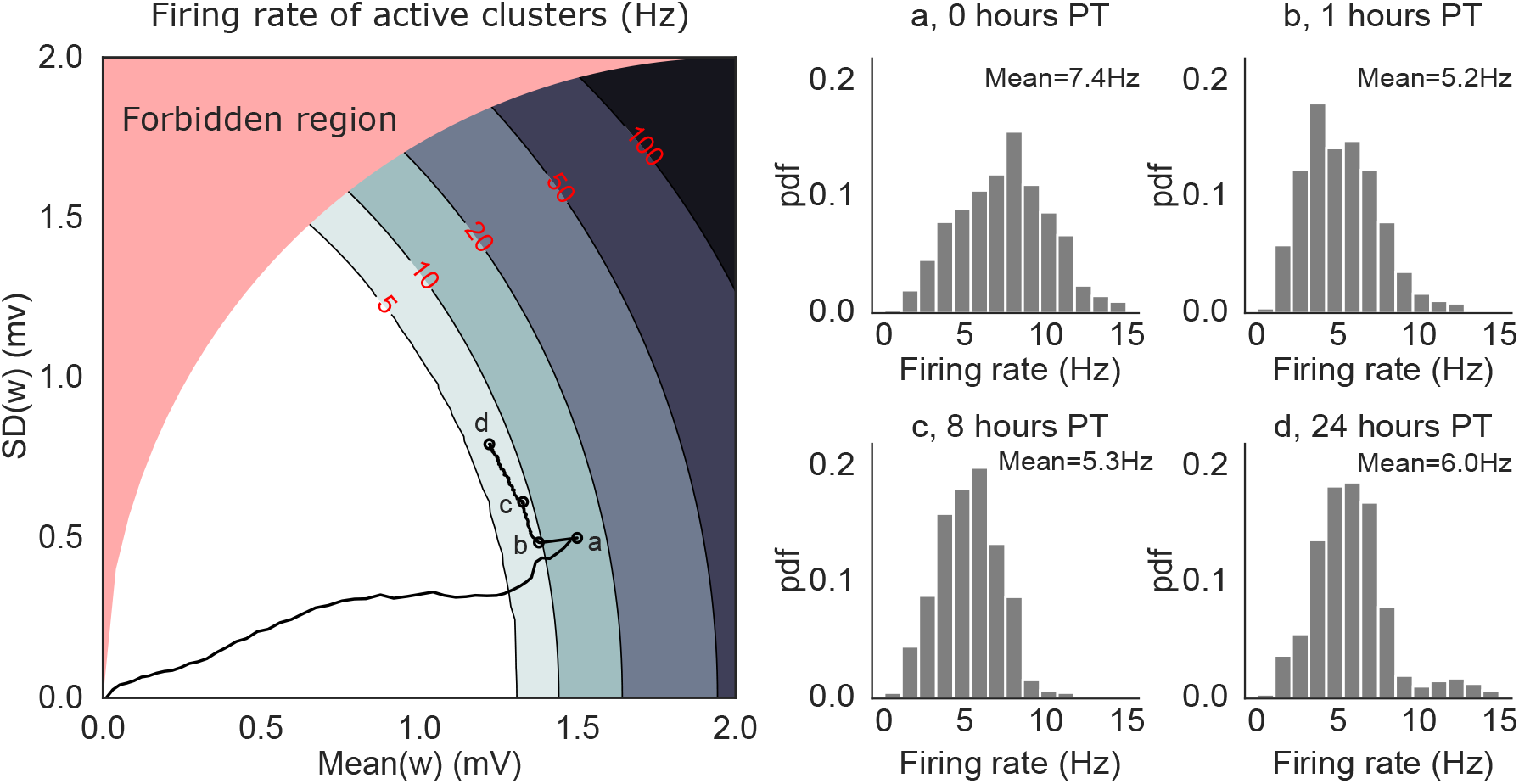
Mean-field solutions to a simplified network model of *N* = 5000 neurons and *Q* = 50 non-overlapping clusters. *Left:* mean field landscape of the network, showing the mean field solutions for the firing rates of active clusters as a function of the mean and standard deviation of the synaptic weights inside clusters. The contour lines mark the mean field firing rates of the active clusters, assuming that at most one cluster can be active. In the white region, all clusters are inactive. The black line shows the trajectory of the average synaptic weights (taken from the simulations shown in Fig. 5) superimposed onto the mean-field landscape, with the circles marked a-d indicating 0, 1, 8 and 24 hours post-training, respectively. *Right:* Normalized histograms (pdf) of single clusters’ firing rates recorded in simulations at time points a-d. The firing rates are in good agreement with the mean field solutions in the left panel except for (a), presumably due to faster dynamics of the synaptic weights compared to the other time points (mean field assumes fixed synapses). The red-shaded region (‘forbidden region’) is not accessible (see Methods for details).

The lowest contour line divides the phases with and without active clusters: this is the only region of the landscape where metastable dynamics is possible; we call it the ‘instability line’ because it separates two regions where neural activity is stable (but note that metastability is possible in a region of finite width around the instability line). Near the instability line, all memories can be quickly reactivated – in fact, they are spontaneously reactivated during metastable dynamics.

The solid black curve in Fig. 6 shows the trajectory of the synaptic weights from a 24-hours network simulation (same simulation as Fig. 5). During training, both the mean and variance of the synapses increase proportionally, leading to a dynamical regime (time point ‘a’) where prolonged (but still transient) activations of single clusters are predicted under mean field (Fig. 5a, ‘plasticity OFF’ raster). However, such prolonged activations are not observed in simulations with plastic synapses (Fig. 5a, ‘plasticity ON’ raster) because they would be terminated by the LTP → LTD transitions. The same mechanism keeps adjusting the synaptic weights after training (Fig. 6 left, a → b) until the network activity and synaptic dynamics are, effectively, in equilibrium. In this regime, metastable dynamics is not the consequence of synaptic fluctuations, but is driven by the network’s own generated fluctuations in neural activity. At this point, switching the plasticity off has no effect on the network dynamics (Fig. 5 left, b-d).

The equilibrium dynamics reached post-training is characterized by very slow changes in the mean synaptic weights – much slower that neural metastable dynamics (Fig. 7a). The equilibrium dynamics is also robust to occasional macroscopic changes that can occur in single synapses, as shown in Fig. 7b, suggesting that memories are supported by the collective behavior of synapses inside clusters. The macroscopic changes observed in some synapses is reminiscent of synaptic volatility (Mongillo et al., 2017), and are presumably due to neural fluctuations in the metastable regime. Our model shows that memories are robust to some degree of synaptic volatility, and that such volatility is a consequence of the interplay between synaptic plasticity and neural dynamics. This interplay keeps the synaptic weights close to the instability line where memories can be quickly reactivated.

**Figure 7:**
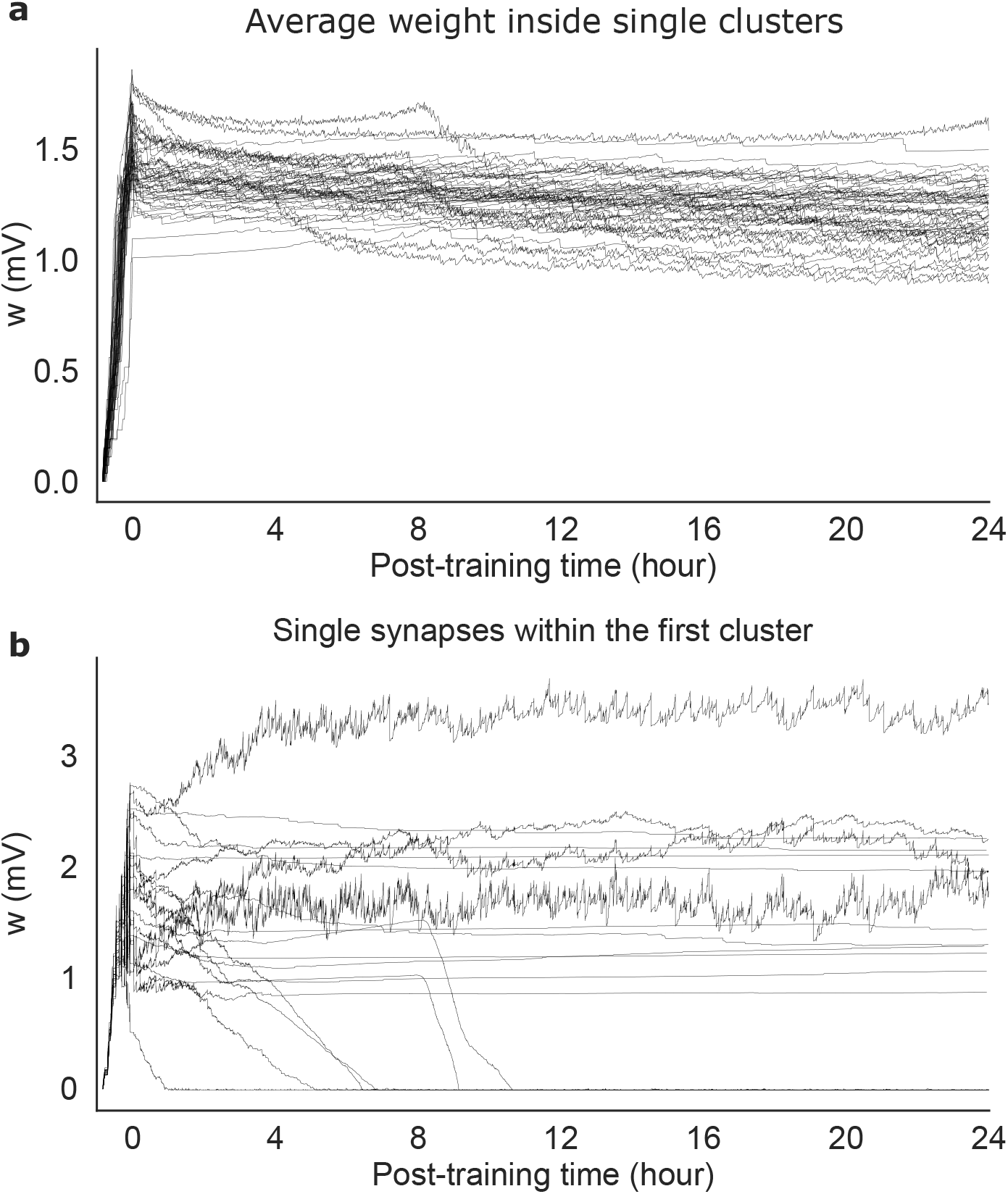
Recordings of single synapses from the network of Fig. 5. (a) Time course of the average synaptic weights within each cluster. (b) Time course of 20 randomly selected synapses within the first cluster. Note that synaptic dynamics is much slower than neural metastable dynamics.

## 3 Discussion

### 3.1 Model features and main results

Metastable neural dynamics, defined as the repeated occupancy of a set of discrete neuronal states occurring at seemingly random transition times (Brinkman et al., 2022), has recently come to the fore as a potential mechanism mediating sensory coding, expectation, attention, navigation, decision making and behavioral accuracy (see (La Camera et al., 2019; Brinkman et al., 2022) for reviews). At the same time, model variations over a basic network of spiking neurons with clustered architecture (Miller and Katz, 2010; Deco and Hugues, 2012; Litwin-Kumar and Doiron, 2012; Mazzucato et al., 2015) have accounted for a wealth of data concerning this metastable dynamics (Mazzucato et al., 2015; Mazzucato et al., 2016; Mazzucato et al., 2019; Lang et al., 2023). One is then led to the following question: how can a cortical circuit be shaped by internal dynamics and externally-driven events so as to converge to the clustered architecture producing metastable dynamics?

The model of synaptic plasticity introduced in this work answers this question by offering a simple yet biologically plausible mechanism capable of cluster formation and metastable dynamics. The plasticity rule builds clusters of neurons with strengthened synaptic connections; after training, the neural activity switches among a number of ‘states’ that can be interpreted as neural representations of the stimuli used for training. Notably, the metastable dynamics generated by learning occurs co-exists, after training, with ongoing synaptic plasticity. This coupled dynamical equilibrium of neural activity and synaptic dynamics is achieved via a self-tuning mechanism that keeps the synaptic weights around the instability line between quiescent and persistent activity (by quiescent activity we mean that no clusters are active). Around the instability line, metastable dynamics results from network ingredients (clustered architecture, quenched random connectivity, recurrent inhibition and the finite size of the network), and although it co-exists with synaptic plasticity, it does not require it. This allows to keep stable representations of learned stimuli in the face of ongoing plasticity, a long-lasting problem known as the stability-plasticity dilemma (Fusi, 2002; Abraham and Robins, 2005; Mermillod et al., 2013; Benna and Fusi, 2016).

Our plasticity rule has several other desirable features. One is biological plausibility, in at least two ways: i) the plasticity rule is local, i.e., it depends only on quantities that are available to the synapse, and ii) it works in networks of spiking neurons (rather than artificial neural networks or firing rate models). Another desirable feature of our rule is that it leads to metastable dynamics with similar properties as those observed in experimental data (Abeles et al., 1995; Jones et al., 2007; Ponce-Alvarez et al., 2012; Mazzucato et al., 2015; Maboudi et al., 2018; Benozzo et al., 2021; Recanatesi et al., 2022; Lang et al., 2023), including an exponential distribution of cluster activation durations with means of few hundreds of ms (Mazzucato et al., 2015). The distribution of synaptic weights after training is itself a slightly skewed unimodal distribution, as observed in real data (Brunel, 2016; Campagnola et al., 2022). Our model also has a minimal number of mechanisms and parameters within a class of similar models that might produce metastable dynamics – for example, a model wherein the presynaptic spike train is replaced by a low-pass filter of it; or a model that imposes an upper bound to the synaptic weights. Those mechanisms are not needed in our plasticity rule. In addition to an adaptive threshold, our model requires only two mechanisms: an attenuation term for LTP (the 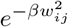 term in Eq. 1) and a spiking term 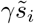 in the equation for the adaptive threshold (Eq. 2). These mechanisms require the introduction of parameters *β* and *γ*, which have a clear biological correlate. *β* controls the amount of LTP. Mechanisms reducing LTP but not LTD have been described *in vitro* by (Delgado et al., 2010), who showed how an increase in total excitatory and inhibitory activity (for example due to cluster activation) can rapidly reduce the amplitude of LTP but not LTD. This mechanism could be mediated through changes in calcium dynamics in dendritic spines and has been proposed as an example of ‘rapid compensatory process’ by (Zenke et al., 2017). The parameter *γ* controls the spiking contribution to the threshold’s dynamics. This term could emerge from cellular mechanisms similar to those responsible for afterhyperpolarization currents (Sah, 1996; Andrade et al., 2012), which are often used in models of firing rate adaptation (Wang, 1998; Ermentrout, 1998; La Camera et al., 2004). Additional compensatory mechanisms, such as inhibitory plasticity or synaptic normalization, were not necessary in our model and therefore were not included.

It is interesting that while the mean synaptic weights inside clusters undergo little change post-training (Fig. 7a), single synapses can undergo macroscopic changes, as shown in Fig. 7b. Changes in synaptic weights (also observed in the absence of learning) has been named ‘synaptic volatility’ and presents a challenge to plasticity models underlying the formation of stable memories (Mongillo et al., 2017). In our model, macroscopic changes in synaptic weights are presumably due to the fluctuations of the neural activity in the metastable regime. Our results show that our model can explain at least some form of synaptic volatility as the consequence of the interplay between synaptic plasticity and neural metastable dynamics. This interplay keeps the synaptic weights close to the instability line where memories can be quickly reactivated.

Another notable feature of our model is its behavior under network scaling. Cortical circuits are variable in size but they typically comprise large numbers of neurons (Shepherd and Grillner, 2010). It is therefore important to test whether models of plasticity maintain their properties as networks are scaled up in size. Scaling up a neural circuit is a necessary operation in several neuroscience theories (e.g., the theory of balanced networks (van Vreeswijk and Sompolinsky, 1998; Renart et al., 2007)) and it can be done in several ways. We have chosen here 2 ways, one that is most intuitive, with order N clusters but fixed cluster size, and one that is most commonly used, wherein both the size and the number of clusters grows as 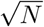 (see (Doiron and Litwin-Kumar, 2014) for a discussion of the different scaling regimes). In both cases, we have found that training leads to the formation of stable clusters generating metastable dynamics, at least for networks up to 20, 000 neurons (larger networks become computationally expensive to simulate). Importantly, both the properties of the dynamics (Fig. S3) and the learned synaptic weights (Fig. 4 and Fig. S4) tend to stabilize as the number of neurons in the circuit grows. In the first scaling regime, we have found that the synaptic decay decreases very fast with *N*, i.e., ∝ 1/*N*.

The behavior of our model under scaling could potentially solve a conceptual problem arising in deterministic network models that try to combine attractor networks with balanced networks (Renart et al., 2007; Doiron and Litwin-Kumar, 2014). These models produce metastable dynamics due to finite size effects, so that in very large networks stable (rather than metastable) cluster activations would prevail (Huang and Doiron, 2017; La Camera et al., 2019; Berlemont and Mongillo, 2022). It is not clear how to obtain metastable dynamics in these models in the limit of very large network size. In the presence of our plasticity mechanism, however, metastable dynamics is the result of a synaptic self-tuning process that keeps the synapses inside the metastable region of the parameter space (Fig. 6). We have directly tested this in a network with 20,000 neurons, with a few thousand synapses per neuron – a reasonable number for neocortex (Shepherd and Grillner, 2010), and we expect that our self-tuning mechanism applies to even larger networks.

### 3.2 Comparison with previous work

Modeling the self-organization of neural circuits via synaptic plasticity has long been an important topic of research (Dayan and Abbott, 2005; Gerstner et al., 2014) and has recently been studied in a number of works bearing various similarities to our work. However, the first successful simulations of spiking networks dynamics with ongoing synaptic plasticity have appeared only about 20 years ago (Del Giudice and Mattia, 2001; Del Giudice et al., 2003; Amit and Mongillo, 2003). Previous models closest to ours are models of voltage-based STDP rules wherein an internal variable (often interpreted as postsynaptic depolarization) is compared to different thresholds for induction of LTP and LTD (Fusi et al., 2000; Clopath et al., 2010; Litwin-Kumar and Doiron, 2014; Zenke et al., 2015). Part of the motivation for some of these models was to reproduce experimental results on both STDP (Markram et al., 1997; Zhang et al., 1998) and the dependence of synaptic plasticity on postsynaptic depolarization (Artola et al., 1990; Ngezahayo et al., 2000; Sjöström et al., 2001). When applied to self-organization of cortical circuits, most previous work in this context has focused on the formation of clusters (often called cell assemblies) for the formation of stable neural activity representing memories. The goal of these works was therefore to obtain stable attractor dynamics after learning. In contrast, the goal of our work was to obtain a stable synaptic matrix that co-exists with a rich, ongoing metastable dynamics.

A notable exception among previous efforts is the model by (Litwin-Kumar and Doiron, 2014). In their model, the authors combined the voltage-based STDP rule of (Clopath et al., 2010) with inhibitory plasticity (Vogels et al., 2011) and synaptic renormalization to obtain metastable dynamics in a network of adaptive EIF neurons. Inhibitory plasticity was used to maintain a target firing rate in the excitatory neurons and prevent the formation of winner-take-all clusters during training. This problem arises due to the fact that, during training, different stimuli target different numbers of neurons creating inhomogeneities in clusters size (Doiron and Litwin-Kumar, 2014) (in our case, the initial inhomogeneity was 10%). We found that in our model, neither inhibitory plasticity nor synaptic normalization was required, presumably thanks to the presence of a dynamic threshold which keeps the synaptic weights bounded as in the BCM rule (Bienenstock et al., 1982). In our model, the dynamic threshold also produces LTD after prolonged cluster activation. This, together with LTP attenuation (the 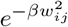 term in Eq. 1; see Fig. S1) helps to prevent the formation of large clusters during training, while also keeping the firing rates within an acceptable range. This confers advantages as it frees up inhibitory plasticity (and possibly other network mechanisms) to accomplish other tasks.

More recently, (Wang and Aljadeff, 2022) have proposed a plasticity rule that produces switching dynamics that resembles ours. In their model, rather than comparing a postsynaptic variable to a threshold for LTP/LTD, a Hebbian term is compared to each threshold. The Hebbian term is the low-pass filter of the product of the low-pass filters of the pre- and post-synaptic spike trains, and was interpreted as a calcium signal responsible for synaptic plasticity (Inglebert et al., 2020). The main goal of (Wang and Aljadeff, 2022) was to present a learning rule resulting in stable and unimodal weight distributions after learning. The authors show that their learning rule has intrinsic homeostatic properties without having to impose an additional homeostatic mechanism on timescales which are much shorter than observed experimentally (Zenke et al., 2017). Perhaps due to these homeostatic properties, they reported metastable dynamics with 2 or 3 stimuli after training. However, as the aim of their work was not to investigate the potential for metastable dynamics, training was only accomplished with few stimuli, and no scaling of the network size was attempted.

We have shown that our plasticity rule allows to learn new stimuli. A similar result was obtained with the plasticity rule of (Litwin-Kumar and Doiron, 2014), which however requires inhibitory plasticity and synaptic renormalization. More recently, (Manz and Memmesheimer, 2023) have proposed a pure STDP plasticity rule that, similarly to our rule, does not require inhibitory plasticity or synaptic renormalization. However, (Manz and Memmesheimer, 2023) focus on the generation of stable clusters producing persistent activity rather than metastable dynamics. Their persistent activity has the signature of fast Poisson noise due to the stochastic nature of their model neurons, but lacks the slow fluctuations characteristic of ongoing metastable dynamics (Litwin-Kumar and Doiron, 2012). We also note that our results on remapping of sensory stimuli is different from the reorganization of clusters described in (Manz and Memmesheimer, 2023), which has been related to representational drift (Ziv et al., 2013; DeNardo et al., 2019; Rule et al., 2019; Masset et al., 2022). In our model, new clusters form due to stimulation by a new set of stimuli rather than drifting of single neuron representations due to the noisy dynamics of plasticity as it occurs in (Manz and Memmesheimer, 2023).

Our model is reminiscent of the BCM rule (Bienenstock et al., 1982), and in fact it could be interpreted as a BCM-like rule for spiking neurons in which the non-linearity of the threshold equation is given by a hyperbolic tangent. More common implementations of BCM utilize a power function (often a quadratic function; see e.g. (Intrator and Cooper, 1992; Dayan and Abbott, 2005)). Another difference with our model is the presence, in the latter, of the additional term 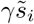 in the threshold dynamics, Eq. 2. In our plasticity rule, this term is required to enforce LTD when the neuron is persistently active (i.e.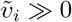), as explained in Sec. 2.1.

### 3.3 Limitations and possible extensions

Our model has a minimal number of ingredients and did not use widespread properties of cortical neurons, such as external noise, firing rate adaptation or short-term plasticity. One reason is model economy, i.e., the desire to use only minimal, essential ingredients. Another reason is that external noise, firing rate adaptation and short-term plasticity all facilitate metastable dynamics (Moreno-Bote et al., 2007; Giugliano et al., 2008; Deco and Hugues, 2012; Cao et al., 2016; Jercog et al., 2017; Setareh et al., 2017; Ballintyn et al., 2019), leading to a possible confound on the role of long-term modifications on generating and maintaining ongoing metastable dynamics.

Although synaptic plasticity depends on pre- and postsynaptic calcium (Sjöström and Nelson, 2002; Shouval et al., 2002; Graupner and Brunel, 2012), we do not consider directly the role of calcium in our model. Rather, our learning rule belongs to the class of voltage-based STDP models (Clopath et al., 2010). These models aim to capture the dependence of LTP and LTD on the membrane potential (Ngezahayo et al., 2000; Sjöström et al., 2001; Wang and Maffei, 2014) while producing STDP curves as an emergent phenomenon. Although voltage-based STDP can capture *in vitro* results where the membrane potential at the electrode is well defined, in a real neuron with an extended geometry the membrane potential is not generally uniform along the dendritic tree. At the time of a postsynaptic spike, one might assume that a back-propagating action potential imposes a uniform voltage on the dendritic tree, at least in proximal dendrites (Stuart and Sakmann, 1994; Nuriya et al., 2006), although faithful propagation depends on many factors including the order of the somatic action potential within a train and the location of the spines (Spruston et al., 1995; Waters et al., 2005). At the moment of a presynaptic spike, however, the voltage at different locations will be different and probably reflect random subthreshold fluctuations. The plasticity rule in such a case would be at the whim of random fluctuations in membrane potential, unrelated to the timing correlation between pre and postsynaptic spikes. We argue that this does not pose a problem in our model because when our postsynaptic variable 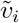 follows random subthreshold fluctuations, our plasticity rule produces no net average synaptic change (Fig. 1b). On the other hand, when the postsynaptic neuron emits a spike, 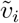 follows closely the transient exponential increase in membrane potential (Fig. 1c), modeling the more uniform effect of the backpropagating action potential along the dendrites. It is even tempting to speculate that, during a prolonged period of active firing, the amplifying buildup of 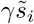 could reflect the facilitation of dendritic calcium spikes by trains of backpropagating action potentials (Larkum et al., 1999).

Our model did not include inhibitory plasticity (Maffei, 2011; Vogels et al., 2013). A benefit of including inhibitory plasticity could be to autonomously set the level of inhibitory activity best suited for metastable dynamics (Zenke et al., 2015). This useful function of inhibitory plasticity contrasts with its homeostatic role in keeping the postsynaptic firing rates close to a desired target value (Vogels et al., 2011; Litwin-Kumar and Doiron, 2014), a role that in our model is fulfilled by the adaptive threshold for excitatory LTP and LTD.

### 3.4 Conclusion

In conclusion, our plasticity rule can successfully structure a network of spiking neurons into an extensive number of clusters (cell assemblies), in a manner that spontaneously generates and maintains metastable ongoing activity. This results from a learning mechanism that keeps the synaptic weights steadily near the threshold for memory reactivation. As a result, one obtains metastable dynamics that coexists with synaptic plasticity. The metastable dynamics supports seemingly random switching among hidden states representing the stimuli used for training, and has several characteristic traits of the metastable dynamics observed in brain regions of rodents and primates. Both metastable dynamics and the learned synaptic structure are stable to random stimulus perturbations, but also flexible enough to be reshaped by new repeated stimuli. Our model could also provide an explanation for the existence of metastable dynamics in large deterministic network of spiking neurons.

## Acknowledgments

We wish to thank Drs. Arianna Maffei, Yonatan Aljadeff and Paul Adams for a critical reading of the manuscript and very useful discussions. This work was partially supported by a U01 grant from the NIH/NINDS Brain Initiative (1UF1NS115779) to G.L.C.

## 4 Methods

### Spiking network model

Here we define the ‘basic’ network, from which all other networks were obtained by scaling up the number of neurons *N* and the number of stimuli *Q* (see ‘Network scaling’ below). The basic network comprised *N*_*E*_ = 800 excitatory and *N*_*I*_ = 200 inhibitory exponential integrate-and-fire (EIF) neurons (described below). The ratio of excitatory to inhibitory neurons was *N*_*E*_/*N*_*I*_ = 4, in agreement with previous studies and experimental observations (Braitenberg and Schüz, 1991; Amit and Brunel, 1997; Shepherd and Grillner, 2010). Before training, neurons were randomly connected with fixed probability (0.2 among excitatory neurons and 0.5 in all the other cases) and constant synaptic efficacies (*w*_*EE*_ = 0.005, *w*_*EI*_ = − 0.34, *w*_*IE*_ = − 0.54 and *w*_*II*_ = − 0.46 mV). Only connected excitatory neurons underwent plasticity.

We modeled the impact of *Q* = 10 random stimuli as follows. Each neuron had a probability (‘coding level’) *f* = 0.1 to be targeted by a stimulus, meaning that the neuron would receive external input during the presentation of that stimulus (we call such neurons ‘responsive’). This resulted in a typical number of 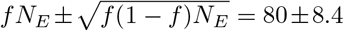 neurons targeted by each stimulus and a mean fraction (1−*f*)^*Q*^ ≈ 0.35 of non-responsive neurons in the basic network (in keeping with similar numbers found in the experimental literature, see e.g. (Perin et al., 2011)). We call a ‘cluster’ a subpopulation of neurons targeted by a given stimulus during training. There were a mean fraction 1 − (1 − *f*)^*Q*^ ≈ 0.65 of neurons in (overlapping) clusters. A randomly picked neuron had a probability *P*_≥2_ = 1 − (1 − *f*)^*Q*^ − *Qf*(1 − *f*)^*Q−*1^ ≈ 0.26 to respond to at least two stimuli, which results in about *P*_*ov*_ = 1 − (1 − *f*)^*Q−*1^ ≈ 0.61 probability for a *responsive* neuron to respond to multiple stimuli (and therefore belong to multiple clusters). Parameter values are summarized in Table 1.

**Table 1:** Model parameters. See the text for details.

| EIF neurons |  |  |
| --- | --- | --- |
| $V_L$ | 0 | leak potential (mV) |
| $V_T$ | 20 | soft spiking threshold (mV) |
| $V_{peak}$ | 25 | peak potential (mV) |
| $V_r$ | 0 | reset potential (mV) |
| $\tau_m$ | 15(E), 10(I) | membrane time constant (ms) |
| $\tau_{syn}$ | 3(E), 2(I) | synaptic time constant (ms) |
| Network (pre-training) |  |  |
| $N$ | from 1,000 to 20,000 | total number of neurons |
| $N_E/N_I$ | 4 | ratio of excitatory to inhibitory neurons |
| $p_{\alpha\beta}$ | $p_{EE} = 0.2, p_{EI} = p_{IE} = p_{II} = 0.5$ | connection probability |
| $w_{EE}$ | 0.005 | $E \rightarrow E$ synaptic efficacy (mV) |
| $w_{EI}$ | -0.34 | $I \rightarrow E$ synaptic efficacy (mV) |
| $w_{IE}$ | 0.54 | $E \rightarrow I$ synaptic efficacy (mV) |
| $w_{II}$ | -0.46 | $I \rightarrow I$ synaptic efficacy (mV) |
| $h^{st}$ | 0.5 | stimulus current ($\text{mV} \cdot \text{ms}^{-1}$) |
| $h^{ext}$ | 1.50(E), 2.19(I) | external current ($\text{mV} \cdot \text{ms}^{-1}$) |
| $Q$ | 10 (basic network) | number of stimuli |
| $f$ | 0.1 (basic network) | coding level |
| Plasticity rule |  |  |
| $\tau_v$ | 50 | time constant of $\tilde{v}_i$ (ms) |
| $\tau_\theta$ | 500 | time constant of $\theta_i$ (ms) |
| $\tau_s$ | 1000 | time constant of $\tilde{s}_i$ (ms) |
| $\theta_a$ | 1 | “amplitude” in Eq. 2 (mV) |
| $g$ | 5 | gain factor in Eq. 2 ($\text{mV}^{-1}$) |
| $\gamma$ | 50 | contribution of spikes in Eq. 2 ($\text{mVms}$) |
| $\beta$ | 0.1 | LTP attenuation factor in Eq. 1 ($\text{mV}^{-2}$) |
| $A_{LTP}$ | 0.005 | LTP strength |
| $A_{LTD}$ | 0.015 | LTD strength |

### Network scaling

For the analysis of Fig. 4, *N, N*_*E*_ and Q were increased with *N*_*E*_ = 0.8*N, Q* = *N*/100 and coding level *f* = 1/*Q*, resulting in a constant mean cluster size of *N*_*Q*_ = *f N*_*E*_ = 80 neurons in all cases. For the analysis of Fig. S4, we used *f* = 1/*Q* with 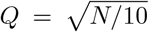 stimuli, resulting in 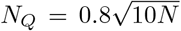 neurons in each cluster (compatible with *QN*_*Q*_ = 0.8*N* = *N*_*E*_ excitatory neurons in total). The factors in the second scaling were chosen so as to agree with the ‘basic’ network for *N* = 1, 000 (*f* = 0.1, *Q* = 10, *N*_*Q*_ = 80).

### Single neuron dynamics

The membrane potential of the *i*-th EIF neuron followed the dynamical equation

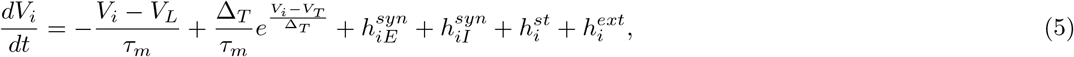

where *V*_*L*_ = 0 mV is the ‘leak’ potential (practically equal to the resting potential in this model), Δ_*T*_ = 1 mV is a ‘sharpness’ parameter related to spike width and upstroke velocity, and *V*_*T*_ = 20 mV is a reference potential somewhat related to the spike threshold (it would be the spike threshold in the limit Δ_*T*_ → 0). A spike was said to be emitted when *V*_*i*_ ≥ *V*_*peak*_ = 25 mV, after which the membrane potential was reset to *V*_*r*_ = 0 mV. The membrane time constant *τ*_*m*_ was 15 ms for excitatory neurons and 10 ms for inhibitory neurons. The term 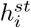 represents a constant stimulus input current which was present only during stimulus presentation. 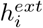 represents a constant external current, presumably coming from more distant regions or other brain areas. The term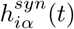, where α ∈ {*E, I*}, is the recurrent synaptic input coming from excitatory and inhibitory neurons of the network, respectively. The synaptic input to neuron *i* obeyed the equation

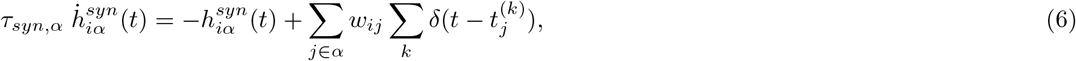

where 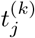 denotes the arriving time of the *k*-th spike of neuron *j, w*_*ij*_ is the synapse weight from neuron *j* to neuron *i*, and *τ*_*syn,α*_ the synaptic time constant of the presynaptic inputs, equal to *τ*_*syn,E*_ = 3 ms for excitatory inputs and *τ*_*syn,I*_ = 2 ms for inhibitory inputs (see Table 1).

### Plasticity rule and training protocol

In our model, only the synapses between excitatory neurons were plastic and they were subject to the following plasticity rule (with *w*_*ij*_ ≥ 0):

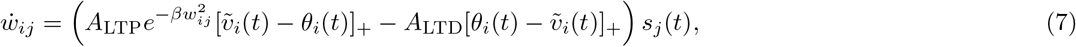

where 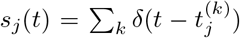 is the presynaptic spike train and [*x*]_+_ := max(*x*, 0) is the rectified linear function. The dynamical threshold *θ*_*i*_(*t*) evolved according to Eq. 2 of the main text:

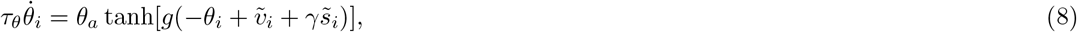

where the notation 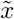 indicates a variable obeying the equation 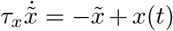, i.e., 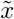 is a low-pass filter of the variable *x*(*t*). Specifically, 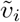 is the exponential voltage term 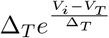 low-pass filtered with time constant *τ*_*v*_ = 50 ms, while 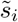 is the postsynaptic spike train low-pass filtered with time constant *τ*_*s*_ = 1 s. *τ*_*θ*_, *θ*_*a*_ and *γ* were constant (see Table 1).

The time-scale of *θ* dynamics is roughly a linear function of its argument (Fig. 1a). Specifically, consider the equation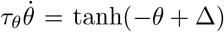, when a change in its fixed point Δ occurs at *t* = 0 (*θ* = 0 when *t* < 0). The time it takes for *θ* to reach half of Δ can be computed to be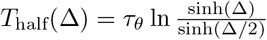, implying a fixed time constant *T*_half_ ≈ *τ*_*θ*_ ln 2 when Δ ≪ 1 and *T*_half_ ≈ *τ*_*θ*_Δ/2 when Δ > 3 (Fig. 1a).

In the basic network, initial training was performed by randomly presenting one stimulus out of *Q* every 2000 ms for a duration of 10 minutes (presentation at random times did not affect the results). Each stimulus lasted 500 ms and and was presented an average of 30 times during the training period. The training time was linearly scaled in larger networks (i.e., 20 minutes for a network with 2*Q* clusters, and so on), resulting in the same mean number of presentations per stimulus in all networks. Perturbing stimuli after training (Fig. 3), whether reoccurring or not, were presented for 200 ms at random times characterized by an exponential inter-event distribution with an average of 10 s.

### Quantification of neural and synaptic activity

A cluster is defined as a subpopulation of excitatory neurons targeted by a given stimulus during training. An active cluster (see below) is interpreted as an internal representation of the corresponding stimulus, and therefore a memory of that stimulus in the absence of external stimulation (active clusters could also represent decision variables and other abstract variables, see e.g. (La Camera et al., 2019)). Cluster activity was measured by the normalized firing rate of its neurons, specifically, for the *q*-th cluster,

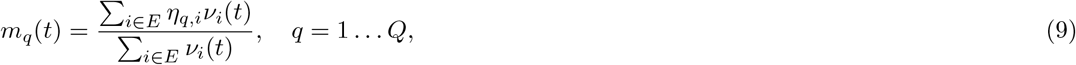

where *ν*_*i*_(*t*) is the firing rate of neuron *i* at time *t* and η_*q,i*_ = 1 if neuron *i* is a target of stimulus *q*, otherwise it is zero. We call *m*_*q*_ the ‘overlap’ between the network activity and stimulus *q*. By definition, 0 ≤ *m*_*q*_ ≤ 1: *m*_*q*_ = 1 implies that only neurons responding to stimulus *q* have non-zero firing rates; *m*_*q*_ = 0 means that the neurons targeted by stimulus *q* are all silent. A memory for stimulus *q* was said to be active if *m*_*q*_ > 0.5. Since ∑_*q*_ *m*_*q*_ ≈ 1, this guarantees that at any given time there is at most one active memory state (∑_*q*_ *m*_*q*_ is not exactly 1 because neurons can be targeted by multiple stimuli).

To quantify the amount of learning, we defined the average synaptic weight among synapses connecting neurons sharing at least one sensory stimulus (dubbed ‘*w*_1_’), and the average weight among neurons that did not share any sensory stimuli (‘*w*_0_’) according to Eq. 3,

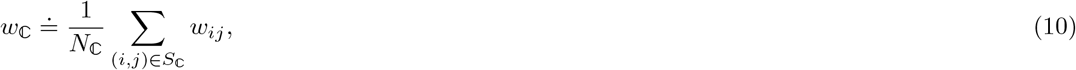

where ℂ ∈ {0, 1}, *N*_ℂ_ is the number of synapses of type ℂ, and *S*_ℂ_ is the set of *ij* indices of synapses of type ℂ. The mean synaptic *change* for synapses of type ℂ, Δ*w*_C_, was defined analogously, i.e., by replacing *w*_*ij*_ with Δ*w*_*ij*_ in Eq. 10:

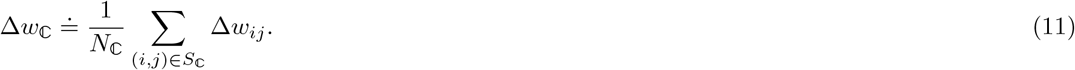

### Scaling laws for synaptic decay rate

Here we derive a qualitative estimate of the synaptic rate of change valid for large N under the scaling used in Fig. 4, i.e., *Q* ∝ *N*_*E*_ ∝ *N* and *f* ∼ 1/*Q* as *N* → ∞. To derive this result, we make a number of assumptions: (i) the presynaptic spike trains are Poisson processes with fixed firing rate that does not depend on *N*; (ii) the post-synaptic term of the learning rule, 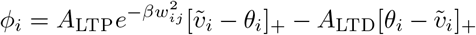, remains finite regardless of *N*; (iii) the initial values (post-training) of the synaptic weights do not depend on *N* (as evident from Fig. 4); (iv) we can temporally separate pre- and postsynaptic terms in the learning rule Eq. 7; and (v) only a finite number of clusters is active an any given time.

Assumptions (i)-(iii) are presumably valid when the cluster size is fixed for different *N* (as is the case in Fig. 4), and are empirically corroborated by our simulations. Assumption (iv) depends on changes in *ϕ*_*i*_ being slow compared to the time scale of single presynaptic spikes (modeled by delta functions). A presynaptic spike *δ* ‘samples’ the value of *ϕ*_*i*_, and since *δ* and *ϕ*_*i*_ have different time scales, they can be treated as approximately independent, especially when the neural activity in active clusters is asynchronous. Assumption (v) seems empirically correct based on our simulations (Fig. 4), in the sense that only a few clusters seem to activate simultaneously.

According to the plasticity rule, Eq. 7, the rate of change in synaptic weight *w*_*ij*_ is proportional to the product of a post-synaptic function of membrane potential, *ϕ*_*i*_, and the presynaptic spike train 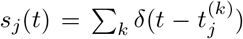. The total synaptic weight change for synapses of a given class ℂ (see Eq. 10) during an interval Δ*t*, is therefore

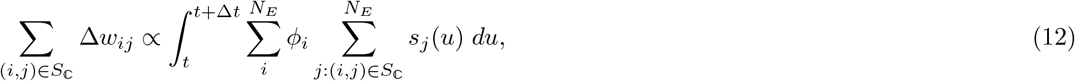

where *i, j* in the right hand side are such that *i, j* ∈ ℂ. The postsynaptic term is the sum of two contributions, one coming from active neurons 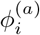 and one coming from inactive neurons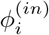, where active neurons are those firing in an active cluster. Of these two contributions, only 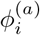 has non-zero mean during learning, because occasional pre- and post-synaptic spikes in inactive neurons produce no average synaptic change, as shown in Fig. 1c.

We further assume that we can separate the pre- and postsynaptic terms in the integral (on account that the latter term is much slower than the former), obtaining

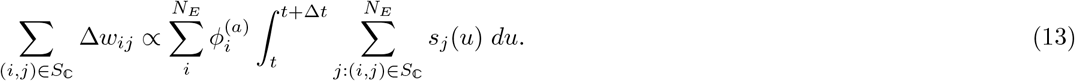

Assuming that only a finite number of clusters is active at any given time, the mean number of active postsynaptic neurons connected by a synapse of type ℂ is proportional to *f N*_*E*_ ∝ *f N* (the mean number of neurons in each cluster), giving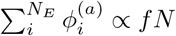.

Assuming Poisson spike trains of average rate ν_ℂ_, the presynaptic term scales as

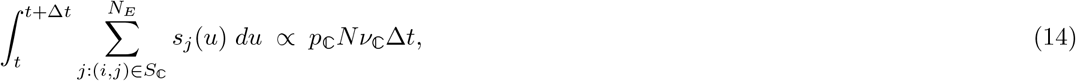

where *p*_ℂ_ is the probability that a synapse is of type ℂ. Putting all together, we get

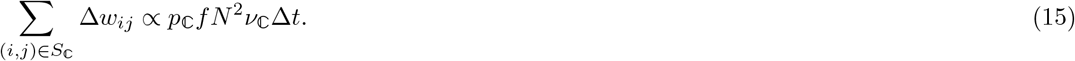

From this equation and Eq. 11 with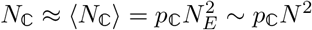, the rate of change of *w*_ℂ_ in an interval Δ*t* is then

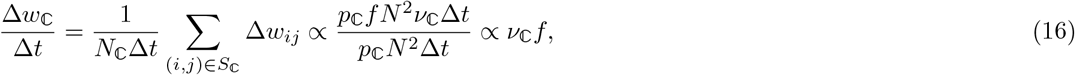

where ν_ℂ_ is the average firing rate of the excitatory presynaptic neurons connected by a synapse of type ℂ. When *f* ∼ 1/*N* (Fig. 4), the cluster size is fixed and the firing rates in the active clusters remain nearly constant with *N* (as empirically found in simulations), leading to Eq. 4 of the main text.

We cannot derive a similar conclusion when 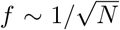 (Fig. S4), since in that case the cluster size increases with *N*. In this case, both the firing rates inside active clusters and the initial synaptic weights post-training change with N in an unknown way.

### Mean-field analysis of network dynamics

For the mean field analysis we considered a simplified network model comprising *N*_*E*_ excitatory and *N*_*I*_ inhibitory exponential integrate-and-fire (EIF) neurons with the same type of random (quenched) connectivity used in simulations. The excitatory population was uniformly partitioned into *Q* clusters: the synapses connecting neurons of the same cluster are denoted by *w*_+_ and those connecting neurons of different clusters are denoted by *w*_*−*_. For simplicity, the clusters were non-overlapping. The *w*_*−*_ synapses were empirically small in simulations, and therefore we fixed them to a constant small value and ignored their dynamics. The *w*_+_ synapses were assumed to be independent samples from a distribution with given mean *µ* and variance *σ*^2^, and were further constrained between 0 and *w*_max_ = 4 mV. This choice of maximum synaptic weight was motivated by the observed distribution of synaptic weights shown in Fig. 5. For a given mean value *µ* ≤ *w*_max_, the standard deviation is bounded by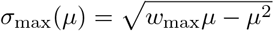, and parameter values above the curve *σ*_max_(*µ*) curve are not allowed (red region in Fig. 6; the bound is saturated by the distribution *P*(*w*_+_ = *w*_max_) = *µ*/*w*_max_, *P*(*w*_+_ = 0) = 1 − *µ*/*w*_max_).

As customary for spiking networks, we performed the mean field analysis under the diffusion approximation, where the input current is characterized by the infinitesimal mean and variance of the synaptic inputs (Amit and Brunel, 1997; La Camera, 2022). The infinitesimal mean and variance to the neurons of the *k*-th excitatory cluster are given by

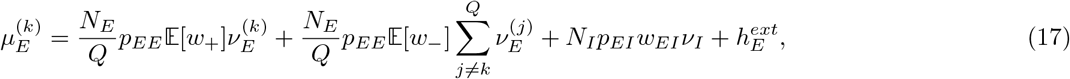

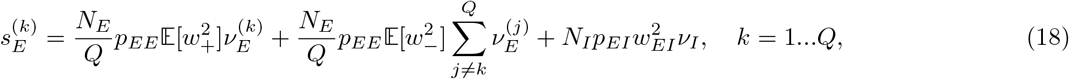

where 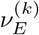 denotes the firing rate of the excitatory neurons in the *k*-th cluster and *ν*_*I*_ is the firing rate of the inhibitory neurons. 𝔼 [·] is the expectation operator. The four terms in the above equations represent the contributions from the *k*-th excitatory cluster, the remaining excitatory clusters, the inhibitory population and the external inputs, respectively. Similarly, the infinitesimal mean and variance of the input to the inhibitory neurons are given by

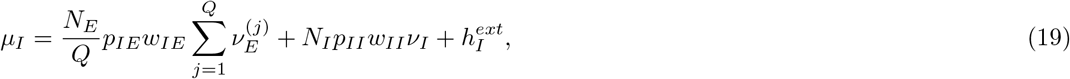

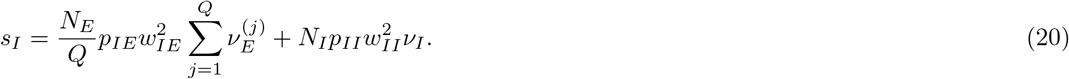

The vector of mean firing rates, 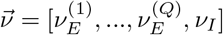 must satisfy the self-consistent equations

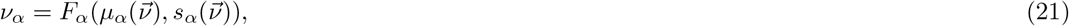

where *F*_*α*_(*µ*_*α*_, *s*_*α*_) is the transfer function of the EIF neuron (Fourcaud-Trocmé et al., 2003). *F*_*α*_ was evaluated numerically by integrating the steady state Fokker-Planck equation describing the EIF model under the diffusion approximation with the algorithm reported in (Richardson, 2007).

## A Supplementary Figures

**Figure S1:**
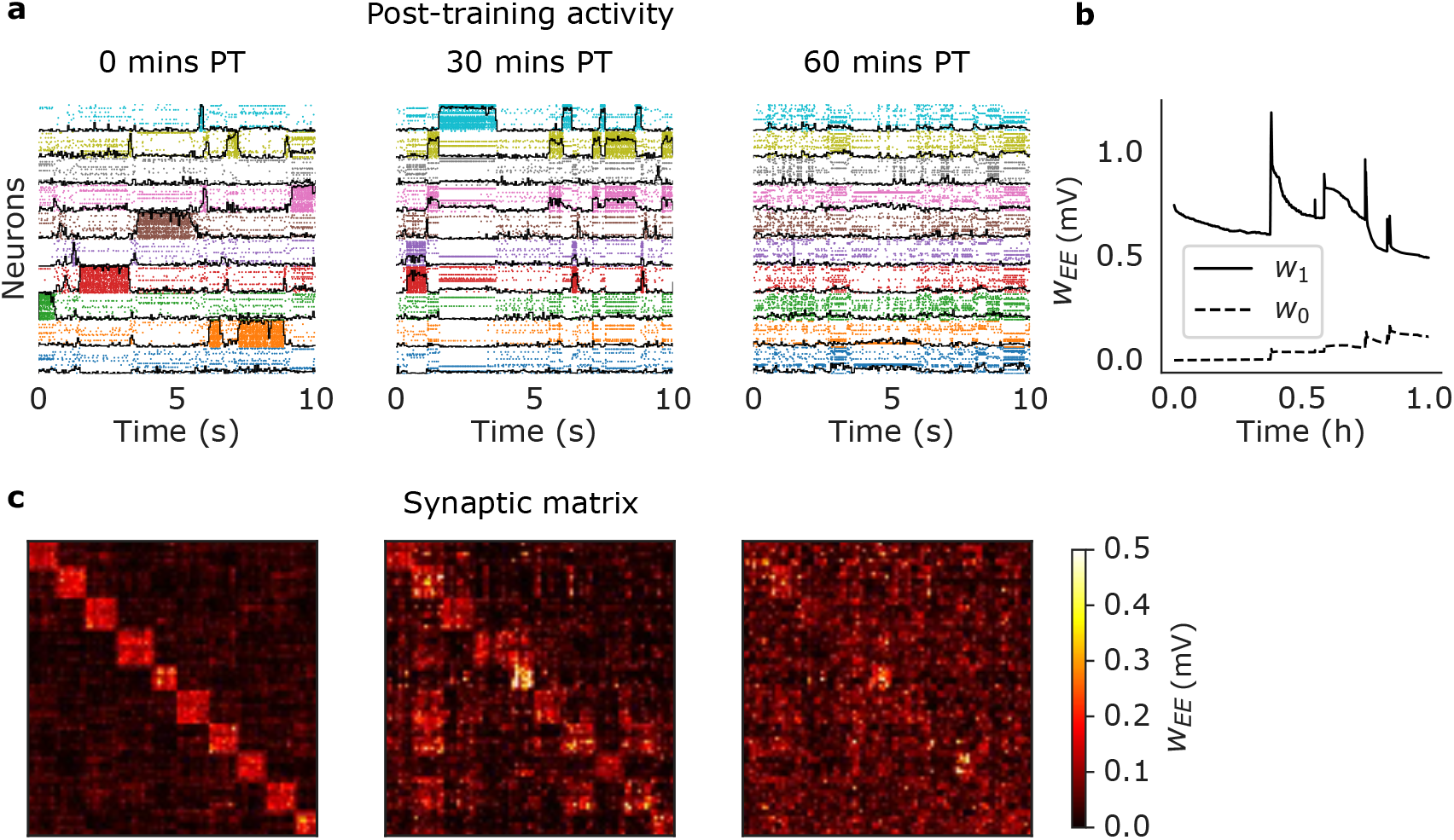
Results from training the basic network of Fig. 2 with *β* = 0 in Eq. 1. (a) Rasterplot of excitatory neurons taken immediately after training (left), 30 minutes after training (middle), and 60 minutes after training (right). Same keys as Fig. 2 of the main text. (b) Averaged post-training excitatory synaptic weights as a function of time. *w*_1_: mean weights across synapses connecting neurons sharing at least one stimulus; *w*_0_: mean weights across synapses connecting neurons sharing no stimuli. (c) Synaptic matrix of the network at the same times as in (a) showing the formation of clusters from the block structure of the matrix.

**Figure S2:**
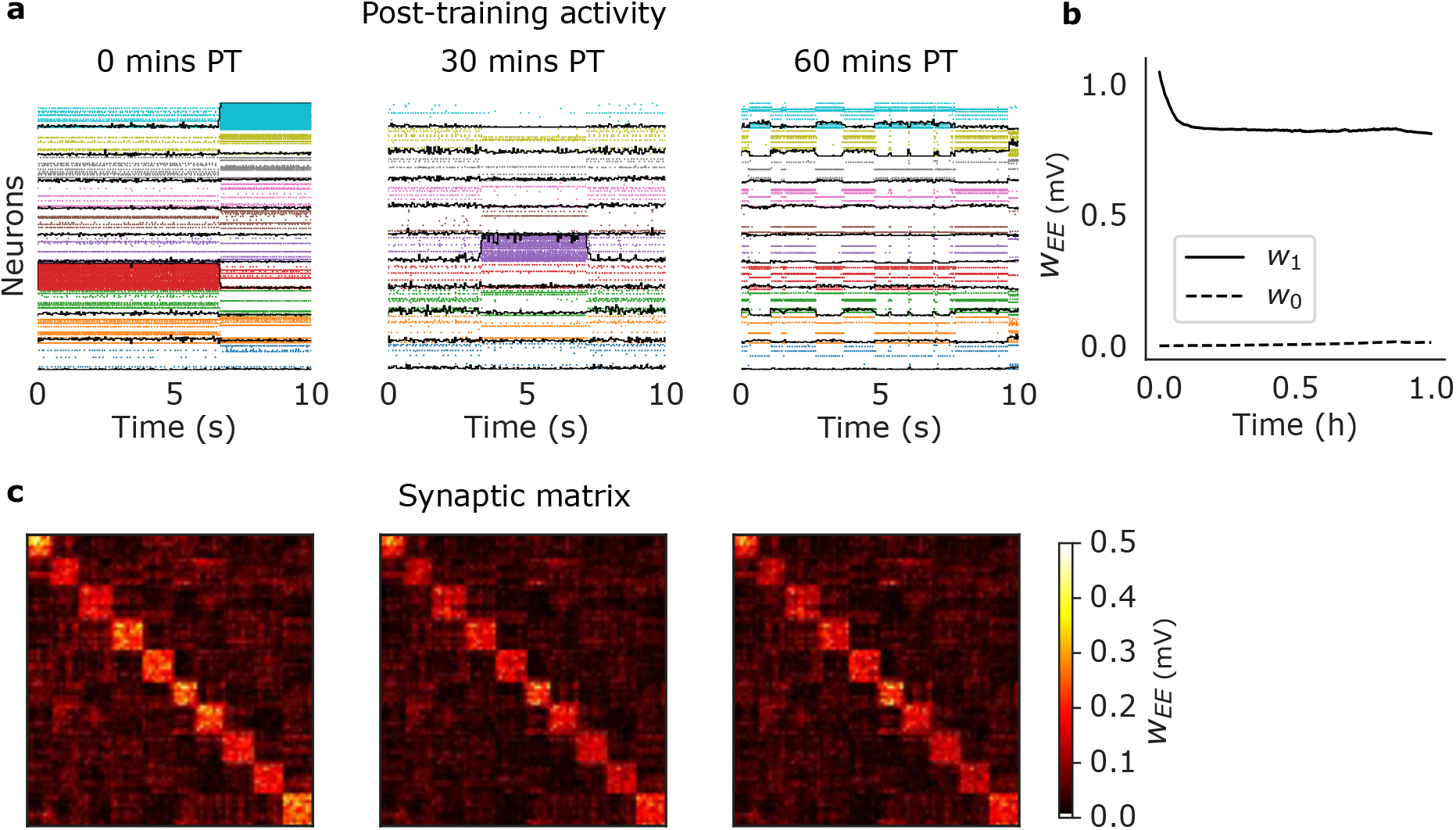
Results from training the basic network of Fig. 2 with *γ* = 0 in Eq. 2. Same keys as Fig. S1.

**Figure S3:**
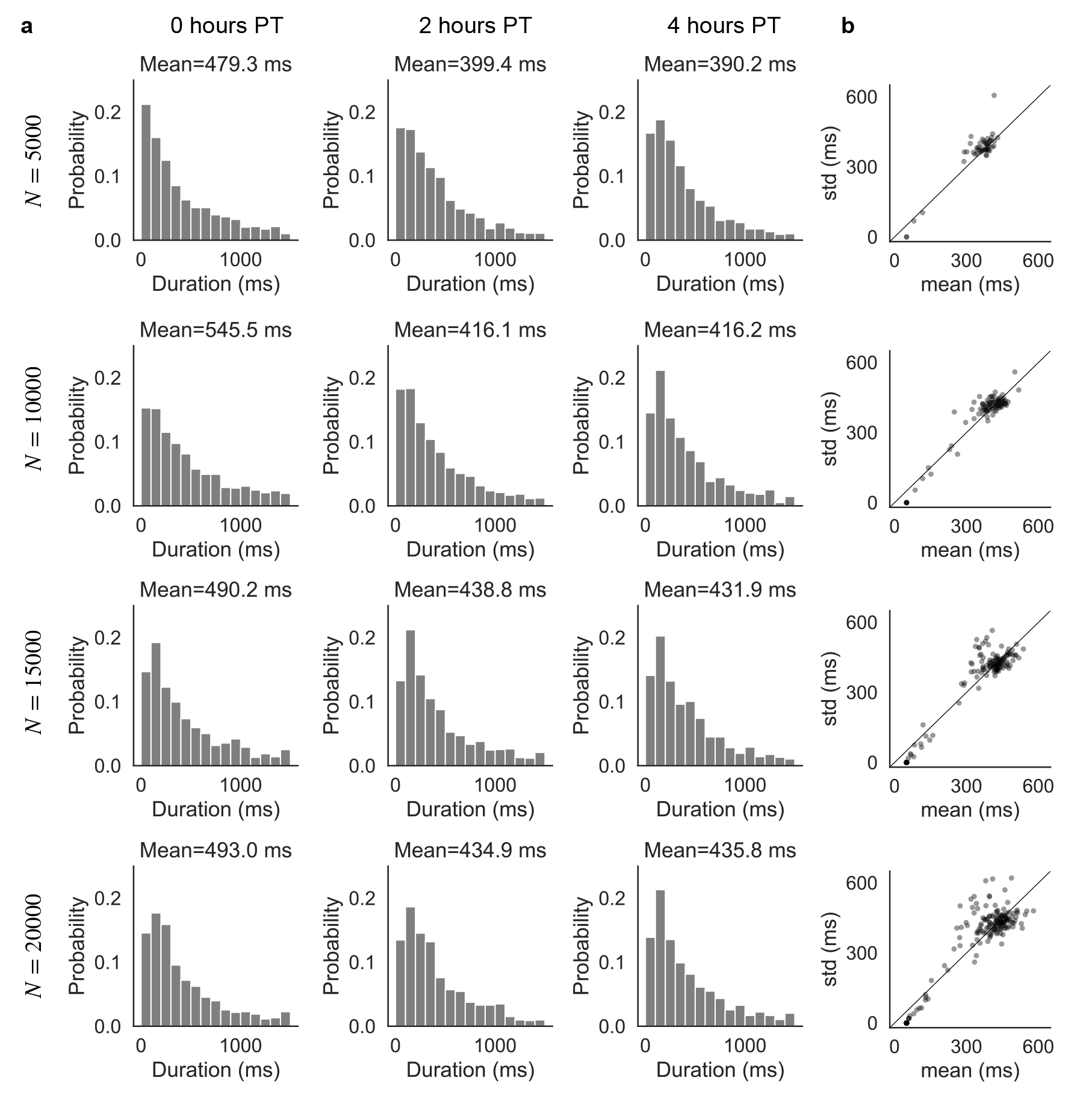
Distributions of durations of cluster activations vs. network size for the networks of Fig. 4. (a) Distributions of durations right after training (left column), 2 hours post-training (middle) and 4 hours post-training (right) for different network sizes. Means tend to decrease with post-training time and increase with network size, approaching stability 4 hours post-training and for *N* ≥ 15, 000. (b) Scatterplots of standard deviation vs. mean of durations for the corresponding networks in (a), superimposed to the identity line. Each circle corresponds to a cluster. For the majority of the clusters, the standard deviations are approximately equal to the means as expected for an exponential distribution.

**Figure S4:**
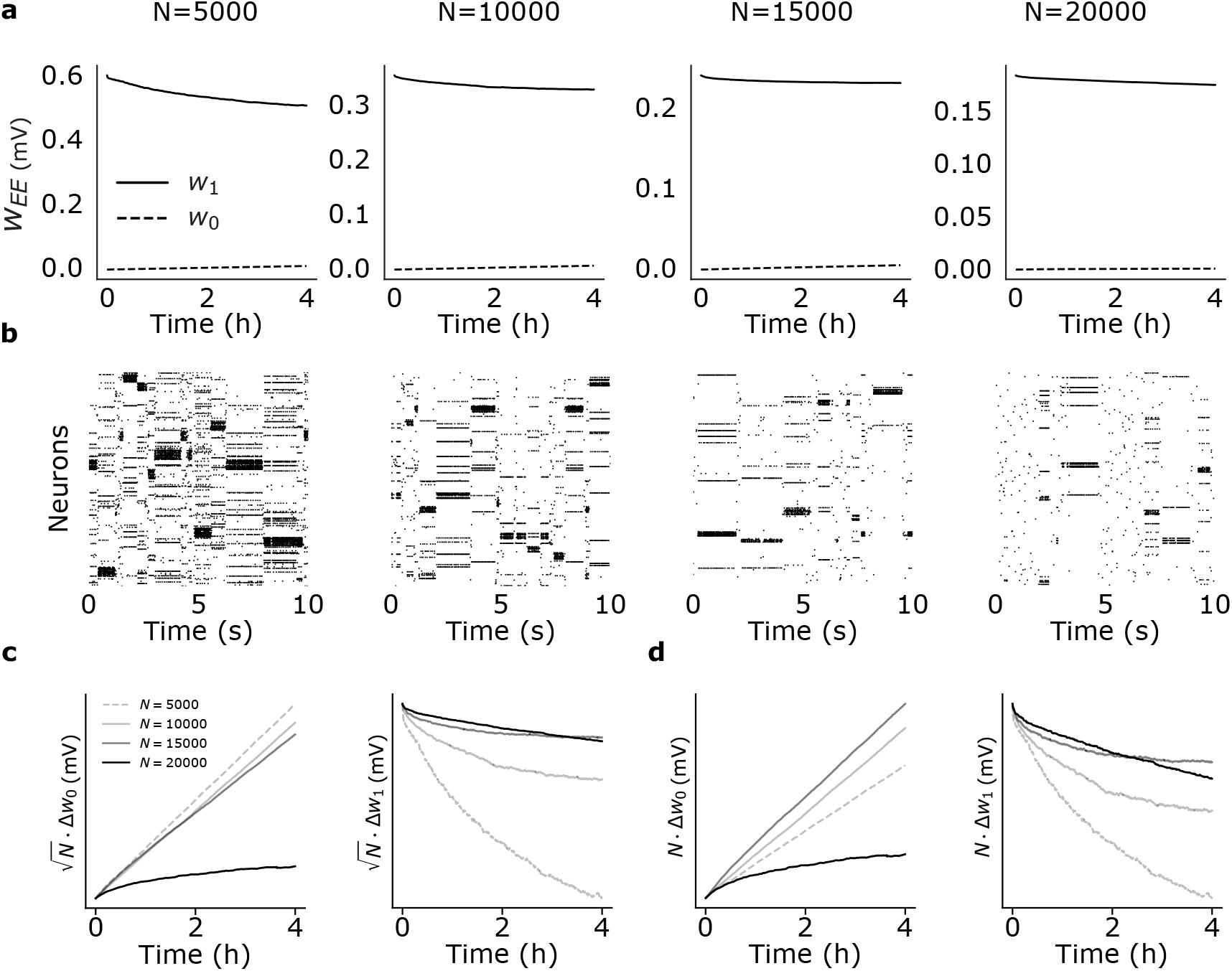
Effect of network size on synaptic dynamics after training. Same as Fig. 4 of the main text but for a different scaling, namely 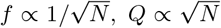 (see the main text). (a) Time dependence of average synaptic weights *w*_1_ and *w*_0_ after training for networks of different size *N*, with *N*_*E*_ = 0.8*N* excitatory neurons. Scaling laws were *f* = 1*/Q* with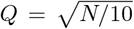, giving 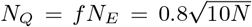 neurons in each cluster (where *Q* and *N*_*Q*_ were rounded to the nearest integer). From left to right, *Q* = 22, 32, 39 and 45. (b) Raster plots of the network’s activity 4 hours after training for the corresponding networks in (a). (c) Plots of 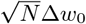 and 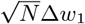 vs. time after training for the different network in (a) (note the difference with panel c of Fig. 4 of the main text). Observations were taken 0 to 4 hours post training. (d) Same as (c) for *N* Δ*w*_ℂ_ vs. time.

